# Validation of the sleep EEG headband ZMax

**DOI:** 10.1101/2023.08.18.553744

**Authors:** Mahdad Jafarzadeh Esfahani, Frederik D. Weber, Merel Boon, Simone Anthes, Tatiana Almazova, Maarten van Hal, Yon Keuren, Carmen Heuvelmans, Eni Simo, Leonore Bovy, Nico Adelhöfer, Milou M. ter Avest, Mathias Perslev, Rob ter Horst, Christiana Harous, Tina Sundelin, John Axelsson, Martin Dresler

## Abstract

Polysomnography (PSG) is the gold standard for recording sleep. However, the standard PSG systems are bulky, expensive, and often confined to lab environments. These systems are also time-consuming in electrode placement and sleep scoring. Such limitations render standard PSG systems less suitable for large-scale or longitudinal studies of sleep. Recent advances in electronics and artificial intelligence enabled ‘wearable’ PSG systems. Here, we present a study aimed at validating the performance of ZMax, a widely-used wearable PSG that includes frontal electroencephalography (EEG) and actigraphy but no submental electromyography (EMG). We analyzed 135 nights with simultaneous ZMax and standard PSG recordings amounting to over 900 hours from four different datasets, and evaluated the performance of the headband’s proprietary automatic sleep scoring (ZLab) alongside our open-source algorithm (DreamentoScorer) in comparison with human sleep scoring. ZLab and DreamentoScorer compared to human scorers with moderate and substantial agreement and Cohen’s kappa scores of 59.61% and 72.18%, respectively. We further analyzed the competence of these algorithms in determining sleep assessment metrics, as well as shedding more lights on the bandpower computation, and morphological analysis of sleep microstructural features between ZMax and standard PSG. Relative bandpower computed by ZMax implied an error of 5.5% (delta), 4.5% (theta), 1.6% (alpha), 0.5% (sigma), 0.8% (beta), and 0.2% (gamma), compared to standard PSG. In addition, the microstructural features detected in ZMax did not represent exactly the same characteristics as in standard PSG. Besides similarities and discrepancies between ZMax and standard PSG, we measured and discussed the technology acceptance rate, feasibility of data collection with ZMax, and highlighted essential factors for utilizing ZMax as a reliable tool for both monitoring and modulating sleep.

## 1. Introduction

Polysomnography (PSG) is the gold standard process of collecting various physiological signals for studying human sleep. Nevertheless, PSG measurement is encumbered by several constraints. A ‘standard’ PSG comprises at least electroencephalography (EEG, ideally from frontal, central and occipital areas), electromyography (EMG, typically submental), and electrooculography (EOG) for scoring sleep stages. The American Academy of Sleep Medicine (AASM, Iber et al., 2007) also recommends including supplementary measurements, e.g., electrocardiography (ECG), blood oxygen saturation level, and body position, while having a standard PSG recording. Most standard PSG systems are costly and cumbersome, confining the majority of sleep studies to the laboratory environment. Furthermore, the process of attaching all standard PSG electrodes is time-consuming and labor-intensive, and the comfort for sleepers wearing multi-channel systems is often poor. Of note, an accurate diagnosis of most chronic sleep abnormalities preferably requires a prolonged period of observation rather than a single night’s examination. However, given standard PSG constraints, it is poorly suited for longitudinal sleep studies. Additionally, apart from the measurement modality itself, the conventional way of scoring sleep data by human experts is also laborious and has an inter-scorer agreement that rarely exceeds 80% (Danker-Hopfe et al., 2009; Rosenberg et al, 2013).

Recent developments in electronics and artificial intelligence (AI) have led to the emergence of various wearable sleep-tracking devices in the consumer market, enabling an alternative to overcome the efforts and discomfort associated with standard PSG procedures. These wearable systems comprise actigraphy (e.g., Ancoli-Israel et al., 2003; Lichstein et al., 2006; Martin et al., 2011; Morgenthaler et al., 2007; Sadeh 2011), smartwatches that fuse data from actigraphy, photoplethysmogram (PPG), and pulse oximeter (e.g., Alfeo et al., 2018; Chang et al., 2018; De Zambotti et al., 2018; Phan et al., 2015; Sun et al., 2017), smart rings with a relatively similar architecture to smartwatches (Altini et al., 2021; Chaudhry et al., 2020; Malakhatka et al., 2021; Mehrabadi et al., 2020; Koskimäki et al., 2018) and importantly, EEG wearables such as headbands that, together with the above mentioned signals, also record scalp EEG (Arnal et al., 2020; Koushik et al., 2019; Mota-Rolim et al., 2019; Onton et al., 2016). Among wearable systems, EEG headbands are the most appropriate candidates to replace standard PSG for longitudinal sleep assessment, given that EEG is needed to provide a direct readout of neural activity including electrophysiological microevents that characterize different sleep stages and are involved in the different biological functions of sleep.

Numerous wearable systems are readily available in the market (see Esfahani et al., 2023 for an overview), despite a paucity of scientific research validating their performance in comparison to standard PSG. Any substitution of the standard PSG with wearables such as EEG headbands first requires validation of their performance. The limited number of EEG channels and the absence of EMG (and occasionally EOG) signals in various wearable headbands pose a challenge for sleep scoring, as these signals are vital for accurate human scoring based on visual inspection of data. Thus, the ‘scorability’ of a wearable is translated into evaluating the robustness of automatic sleep scoring (autoscoring) algorithms. Validation studies should also focus on determining subjective and objective sleep quality parameters and encompass comprehensive assessments such as analyzing signal power across different frequency bands and determining the characteristics of microstructural features within different stages of sleep.

In recent years, several (commercial) EEG wearables have been referenced in the literature, including Dreem (Dreem, Paris, France), SleepLoop (Mobile Health Systems Lab, Zurich, Switzerland), and ZMax (Hypnodyne Corp., Sofia, Bulgaria). The Dreem headband is an example of an ambulatory sleep tracker that employs five EEG channels (F7, FpZ, F8, O1, O2), an accelerometer, and a pulse oximeter, and its performance has been scientifically validated. Arnal et al. (2020) evaluated the performance of the Dreem headband compared with a standard PSG in terms of EEG signal quality, accuracy of the heart rate and breathing rate estimation, and the performance of the autoscoring. In this study, the researchers found slight differences in the relative spectral power estimation of EEG signals between the Dreem headband and PSG, i.e., mean percentage error of 15 ± 3.5%, 16 ± 4.3%, 16 ± 6.1%, and 10 ± 1.4% for α, β, λ, and θ bands, respectively. The automatic sleep scoring algorithm of Dreem indicated acceptable outcomes when compared to the consensus of five professional human scorers (F1-score: 83.8 ± 6.3 % in autoscoring vs. 86.3 ± 7.4 % in the consensus of human scorers).

SleepLoop is a portable sleep-tracking system that offers eight configurable ExG channels which might be used as EEG, EOG, and EMG. This headband was shown to have a comparable signal quality with respect to standard PSG, reaching correlation values of 0.98 and 0.99 for the delta and sigma frequency bands, respectively (Ferster et al., 2019). The evaluation of SleepLoop has not been limited to sleep tracking only; the researchers also investigated its performance in real-time to apply closed-loop auditory stimulation (CLAS) in healthy elderly and Parkinson’s disease patients and attained comparable outcomes with intensive in-lab systems (Ferster et al., 2022).

Another EEG wearable that has been widely used in sleep studies is the ZMax EEG headband. Despite the broad utilization of ZMax for sleep monitoring, non-rapid eye movement (non-REM) sleep modulation through CLAS and targeted memory reactivation (TMR), and rapid eye movement (REM) sleep modulation (Berger, 2022; Bradshaw 2019; Esfahani et al., 2022b; Mota-Rolim 2018; Stocks et al., 2020; Talamini et al., 2022; Van Trigt et al., 2022), scientific validation and benchmarking of ZMax with respect to standard PSG (further denoted as PSG) has yet to be performed.

Among the available EEG wearables are some such as Dreem that utilize dry electrodes, whereas SleepLoop and ZMax use disposable wet electrodes. Wet electrodes typically exhibit lower impedance and thus an enhanced signal quality; however, they come at the cost of replacing the electrode patch after a few uses, incurring extra costs. Furthermore, while the dry electrodes generally have a static positioning, wet electrodes may be repositioned to a different spot based on the research goals. Also, the wet electrodes generally represent superior skin contact when compared to the dry electrodes, which decreases the likelihood of electrode detachment during the night. Headbands which provide ExG channels are highly advantageous as they enable customization of the required montage according to research objectives, e.g., employing a few EMG channels when studying REM sleep. SleepLoop is one of the few headbands with this feature, whereas the majority of the headbands do not contain EMG recording. Enabling cloud computing or real-time data transmission to a computer as in ZMax is another intriguing feature of the wearables that provide extensive computational power through pc resources. This feature assists with developing desired software for various purposes, such as for online detection of a specific sleep oscillation and subsequently modulating it. We believe that ZMax stands among the wearables which employ a reasonable number of the above mentioned features, rendering it a proper option for longitudinal sleep monitoring and modulation.

In this study, we investigated five different datasets, four of which were collected with simultaneous ZMax-PSG recordings from healthy participants either at home or in a laboratory setting. We compared the performance of the autoscoring algorithm of the ZMax headband, i.e., *ZLab*, and a new open-source autoscoring algorithm, dubbed *DreamentoScorer*, with the consensus of human scorers based on PSG as the ground truth. The sleep assessment metrics were computed by each autoscoring approach and compared with the results derived from the ground truth scorings. We analyzed the relative bandpower of the signals to evaluate the agreement between the measurement modalities. The possibility of detecting microstructural features such as SOs and spindles during non-rapid eye movement (non-REM) and rapid eye movement (REM) events during REM sleep has been explored, demonstrating the corresponding agreement between measurement modalities using Bland-Altman plots. We also examined the influence of wearing ZMax EEG headband on subjective outcomes such as sleep quality, morning mood, number of awakenings, as well as its comfort and disturbance level. The feasibility of data collection with ZMax has been estimated as the percentage of useful data. Additionally, an assessment of the feasibility of gathering and analyzing extensive longitudinal data using ZMax within a ‘patient’ target group was conducted. Finally, we discussed the findings regarding the discrepancies and similarities of ZMax outcome with respect to PSG, as well as potential future directions.

## 2. Methods

An overview of the analysis pipeline of this study is provided in Figure 1. All the PSG data from different datasets (datasets 1-4) were manually scored by human experts according to the AASM protocols and then the consensus was made for datasets comprising more than one scorer to serve as the ground truth. Our behavioral analysis was focused on dataset 2, which was specifically designed for the validation of ZMax EEG headband and thus included relevant questionnaires. For our analysis, we applied pre-processing steps such as bandpass filtering in the primary sleep frequency range of 0.3 - 30 Hz, resampling the PSG data to the same sampling frequency as ZMax, and then aligning the PSG and ZMax epochs based on predefined tasks, e.g., eye movements prior to sleep. Following the pre-processing steps, the main analysis was conducted, including training and evaluating the performance of our open-source autoscoring model, i.e., DreamentoScorer, as well as testing the ZLab autoscoring algorithm, comparing sleep statistics determined using different methods, time-frequency representation (TFR) comparison, bandpower assessment, and an in-depth analysis on determining the characteristics of sleep microstructural features using ZMax and PSG.

**Figure 1.**
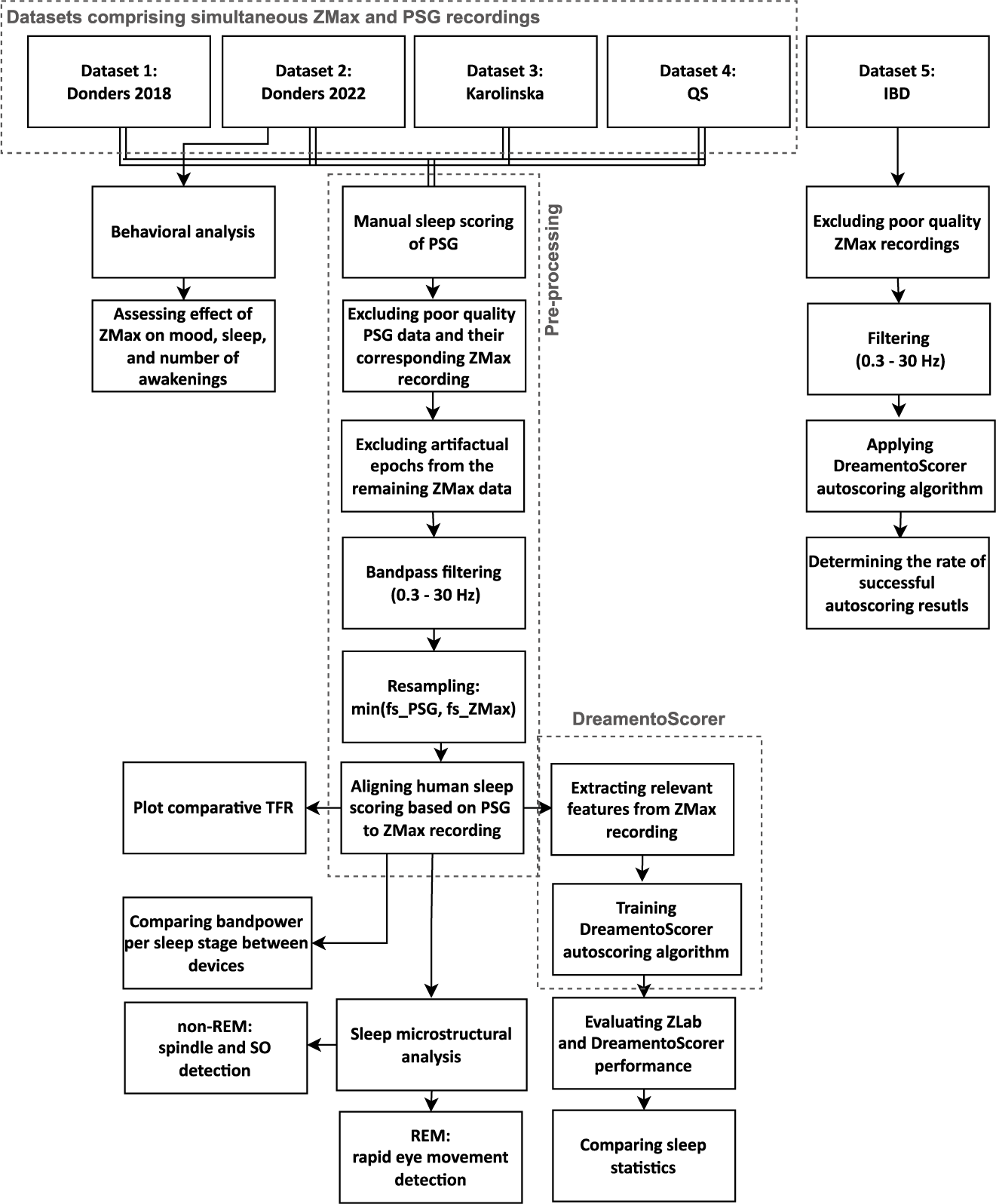
Analysis pipeline of the study. Datasets 1-4 encompassed simultaneous ZMax-PSG recordings, whereas dataset 5 only included ZMax recordings. fs_PSG: sampling frequency of PSG (Hz), fs_ZMax: sampling frequency of ZMax (Hz), PSG: polysomnography, REM: rapid eye movement, SO: slow-oscillation, TFR: time-frequency representation.

### 2.1. Analysis layers

Datasets 1-4 have all undergone three different analysis layers, namely: (1) (auto-)scoring and sleep metrics assessment, (2) signal quality and non-REM features analysis, and (3) REM features analysis. Therefore, different inclusion criteria have been established, as analysis layers two and three required specific EEG montage. For the (auto-)scoring assessment (layer 1), the criteria required a minimum of

∼75% ‘scorable’ data from both ZMax and PSG, with the overall recording duration being at least (approximately) 5.5 hours. This means that there was no specific requirement for the quality of individual channels from the PSG; instead, the combined data from all available channels needed to be acceptable for the human scorer to score at least 75% of the data. On the other hand, for the assessment of signal quality and non-REM features analysis (analysis layer 2), which relied on the closest channel from PSG in relation to ZMax, different conditions were applied. In addition to data scorability (layer 1 criteria), layer 2 mandated that the F3 and F4 channels from the PSG had to meet acceptable quality standards (see also section 2.5. Preprocessing). For the third analysis layer, in addition to layer 1 criteria, we considered the quality of horizontal EOG channels from the PSG, while no specific requirements were imposed on the other channels. Consequently, the number of included data for each analysis layer may not be identical (see Table 1 for an overview).

**Table 1.** Dataset overview. Analysis layer 1: (auto-) scoring and sleep metrics assessment. Analysis layer 2: signal quality assessment in addition to non-REM features analysis based on frontal channels (F3 and F4 from the PSG). Analysis layer 3: REM features analysis using dedicated EOG channels from the PSG. Exclusion criteria correspond to analysis layer 1 as the primary step. Slight differences in exclusion rates of analysis layer 1 with respect to both 2 and 3 are related to data quality of F3 and F4, and EOG channels of PSG, respectively. ^✞^: Datasets 1, 2, and 4 have been collected in Donders institute, however, with completely different researchers and experimental teams as well as instruction procedure for participants. *: In dataset 5, 366/512 (71%) met the inclusion for analysis layer 1, i.e, at least 5.5 hours in duration while including at least 75% scorable data, but given the absence of PSG, we only used completely clean data which did not require any specific artifact rejection step which resulted in an inclusion rate of 280 / 512 (55%).**: The first three nights collected from the first participant showed irregular sleep microstructural patterns (probably due to electrodes misplacement/software configuration and despite having clean signal quality throughout the recording, the data were excluded based on human scorer suggestion. ***: 17/25 data with poor quality were collected during the last 18 measurements indicating an internal issue with the headband.

|  | **Dataset 1:**<br>**Donders 2018** | **Dataset 2:**<br>**Donders 2022** | **Dataset 3:**<br>**Stockholm Uni.** | **Dataset 4: QS** | **Dataset 5:**<br>**MindIBD** |
| --- | --- | --- | --- | --- | --- |
| **N recruited Participants** | Cohort 1: 11<br>Cohort 2: 12 | 37 | 26 | 1 | 142 |
| **Age (mean ± SD) years** | Cohort 1: 21.1 ± 3.4<br>Cohort 2: 23.8 ± 3.6 | 21.5 ± 3.7 | 27.8 ± 6.1 | 35 | 48.62 ± 13.96 |
| **Gender** | Cohort 1: 4/11 (36 %) females<br>Cohort 2: 9/12 (75 %) females | 25/37 females (68 %) | 14/23 females (54 %) | 1 male | 91 females (64 %) |
| **N available parallel recordings** | 23 | 97 | 57 | 68 | N/A |
| **N inclusion for analysis layer 1** | 9 / 34 (26 %) | 66 / 96 (69 %) | 17 / 57 (30 %) | 43 / 68 (63 %) | 280 / 512 (55%)* |
| **N inclusion for analysis layer 2** | 6 / 23 (26 %) | 61 / 96 (64 %) | 13 / 57 (23 %) | 38 / 68 (56 %) | N/A |
| **N inclusion for analysis layer 3** | 8 / 23 (35 %) | 63 / 96 (66 %) | 14 / 57 (25 %) | 41 / 68 (60 %) | N/A |
| **Main reason for exclusion** | - Entirely noisy ZMax: 3<br>- Entirely noisy PSG: 3<br>- <75% useful data: 18<br>- < 5.5 hours, but clean: 1 | - Entirely noisy ZMax: 1<br>- Entirely noisy PSG: 3**<br>- <75% useful data: 19<br>- < 5.5 hours, but clean: 2 | - Entirely noisy ZMax: 17<br>- Entirely noisy PSG: 0<br>- <75% useful data: 22<br>- < 5.5 hours, but clean: 1 | - Entirely noisy ZMax: 3<br>- Entirely noisy PSG: 2<br>- <75% useful data: 20 ***<br>- < 5.5 hours, but clean: 0 | N/A |
| **Study environment** | Home | Home | Lab | Home | Home |
| **Research Institute** | Donders <sup>#</sup><br>Institute for Brain, Behaviour and Cognition – research team 1 | Donders Institute for Brain, Behaviour and Cognition – research team 2 | Stockholm University | Donders Institute for Brain, Behaviour and Cognition – research team 3 | Four Dutch hospitals in Nijmegen (2), Arnhem and 's-Hertogenbosch |
| **PSG system** | SOMNOscreen | SOMNOscreen | Vitaport | SOMNOscreen | N/A |
| **Available EEG channels** | Frontal (F3, F4), Central (C3, C4), Occipital (O1, O2) | Frontal (F3, F4), Central (C3, C4), Occipital (O1, O2) | Frontal (F4), Central (C3, C4), Occipital (O2) | Frontal (F3, F4), Central (C3, C4), Occipital (O1, O2) | N/A |

The aim of the analysis layers 2 and 3 was to compare the signal quality between ZMax and PSG. Thus, to remove the impact of autoscoring performance (analysis layer 1) on these layers, we employed human scoring from the PSG and aligned it with ZMax, rather than using the autoscoring results. This way, we have ensured the consistency in the scoring across measurement modalities, which allowed us to solely compare the signals and evaluate their similarities as well as differences in terms of the outcome measures.

### 2.2. Measurement modalities

The Lite version of ZMax EEG headband comprises two frontal EEG channels (F7-Fpz, F8-Fpz), a tri-axial accelerometer, and a PPG sensor, all with a sampling frequency of 256 Hz. The advanced version in addition includes an ambient light sensor, a microphone (to measure ambient noise), a thermometer, and nasal/SpO2 sensors. Intermediate versions with different kinds of sleep modulation functionalities are also available. The sleep EEG recording functionality tested in the present study is the same for all versions of the headband. For any version, a hydrogel electrode patch has to be mounted on the ZMax headband for EEG recordings. The disposable patches can be used for multiple nights.

The PSG system in datasets 1, 2, and 4, was SOMNOscreen™ (Somnomedics GmbH, Randersacker, Germany), whereas dataset 3 used the Vitaport system (TEMEC Instruments, Kerkrade, Netherlands). SOMNOscreen is a portable PSG system comprising six EEG channels (F3, F4, C3, C4, O1, O2) with a mutual reference to Cz and a ground on Fz and can be further referenced to the Mastoid channels. It also includes two horizontal EOG, three chin EMG, and ECG channels. Recordings through the Vitaport system involve four EEG channels (F4, C3, C4, O2) with a mutual reference to Cz and a ground on Fz and can be referenced to the Mastoid channels. Recordings also include two horizontal EOG, a single chin EMG and ECG signals.

### 2.3. Datasets

This study encompasses five datasets. The first four datasets include (1) Donders 2018, (2) Donders 2022, (3) Stockholm University, and (4) Quantified self (QS) and comprise participants’ nocturnal sleep simultaneously recorded using ZMax and PSG from healthy populations either at home or in lab environments, whereas dataset (5) MindIBD includes nocturnal sleep recordings only through ZMax from patients with inflammatory bowel disease (IBD) in home settings.

1. **Donders 2018:** This dataset comprised two cohorts, cohort 1 with N = 11 (4 females, mean age 21.1 ± 3.4 years std) and cohort 2 including N = 12 participants (9 females, mean age 23.8 years ± 3.6 std). Both groups underwent three nights of nocturnal data collection using ZMax in a home setting. The first group experienced a single night of PSG recording in parallel with ZMax, whereas in the second group, there were two nights of data collection with both systems, one while ZMax was bridged to be recorded through PSG and the other with parallel recording. For analysis in the current investigation, only the parallel recordings were considered. Out of 34 simultaneous ZMax-PSG data 23 included ‘parallel’ recordings, among which 9, 6, and 8 recordings met the inclusion criteria for analysis layers 1, 2, and 3, respectively.
2. **Donders 2022:** a total of N = 37 healthy young participants were included in the study; of which N = 32 completed the study (22 females, mean age 21.59 years ± 3.78 std). The overall duration of the study was six weeks (see Figure 2), starting with an intake session, followed by the pre-experimental period (weeks 1 and 2), continuing towards the experimental period (weeks 3, and 4), and ending with the post-experimental period (weeks 5, and 6). Participants completed baseline questionnaires at the intake and then for the following 43 consecutive days completed daily dream diaries and questionnaires regarding sleep quality. During the experimental period, nocturnal sleep was recorded every night with ZMax and during the first, 8th, and 15th nights simultaneously with the PSG. Before the first experimental night, participants received a video together with a brochure containing the instructions on how to wear, maintain, and record data with ZMax and the PSG. The PSG systems was placed in the lab, and the participants started recording themselves at home. The participants practiced this procedure with the experimenters on the first experimental night, before receiving the PSG and leaving the lab to sleep at home. From the 97 simultaneous ZMax and PSG recordings, 66, 61, and 63 met the inclusion criteria of analysis layers 1, 2, and 3, respectively.
3. **Stockholm University:** N = 23 participants (14 females, mean age 27.82 years ± 6.07 participated in the study. Participants spent three non-consecutive nights in the sleep lab, i.e., a habituation night and two experimental nights. Each night, participants went to bed at their habitual bedtime (ranging from 21:30 to 00:00) and awoke at their habitual wake time (ranging from 04:00 to 07:30). On one of the experimental nights, participants snoozed in bed the last 30 minutes before their habitual wake time. Each night, participants wore several actigraphy monitors on their body, along with the ZMax headband and PSG (Vitaport system). Participants also completed a motivation questionnaire each evening before bed. Upon awakening, participants also produced two saliva samples, completed a sleep questionnaire and completed a cognitive test battery on a smartphone. Participants completed three more cognitive test batteries throughout the day (directly upon awakening; 40 minutes after awakening, around 12:30 pm and around 3:30 pm). From the 23 participants, 17, 13, and 14 recordings were included for analysis layers 1, 2, and 3, respectively.
4. **QS:** the QS dataset consists of various physiological measurements from a citizen neuroscientist (N = 1, male, aged 35 years) over the course of 34 months (Sikder et al, 2022; ter Horst & Dresler 2022). Nocturnal sleep was recorded every night with ZMax, whereas an additional PSG was mounted on the subject once per week to record nocturnal sleep simultaneously. From the 96 nights with simultaneous ZMax and PSG recordings, 43, 38, and 41 were included in analysis layers 1,2 and 3, respectively.
5. **MindIBD:** The MindIBD trial is a multicenter randomized controlled trial that investigates the effectiveness of Mindfulness-Based Cognitive Therapy (MBCT) in patients with IBD (ter Avest et al., 2023). The trial primarily focuses on the effectiveness of MBCT in reducing psychological distress in IBD patients, but the effects on sleep quality, fatigue, disease activity and -control, and quality of life are also outcomes of interest. Within this trial, a targeted number of 136 patients from four Dutch hospitals with a confirmed IBD diagnosis in remission and suffering from psychological distress (Hospital Anxiety and Depression Scale total score ≥ 11) are allocated 1:1 to MBCT + treatment as usual or treatment as usual alone. Participants were asked to sleep with ZMax EEG headband for three consecutive nights, at both baseline and post-intervention.

**Figure 2.**
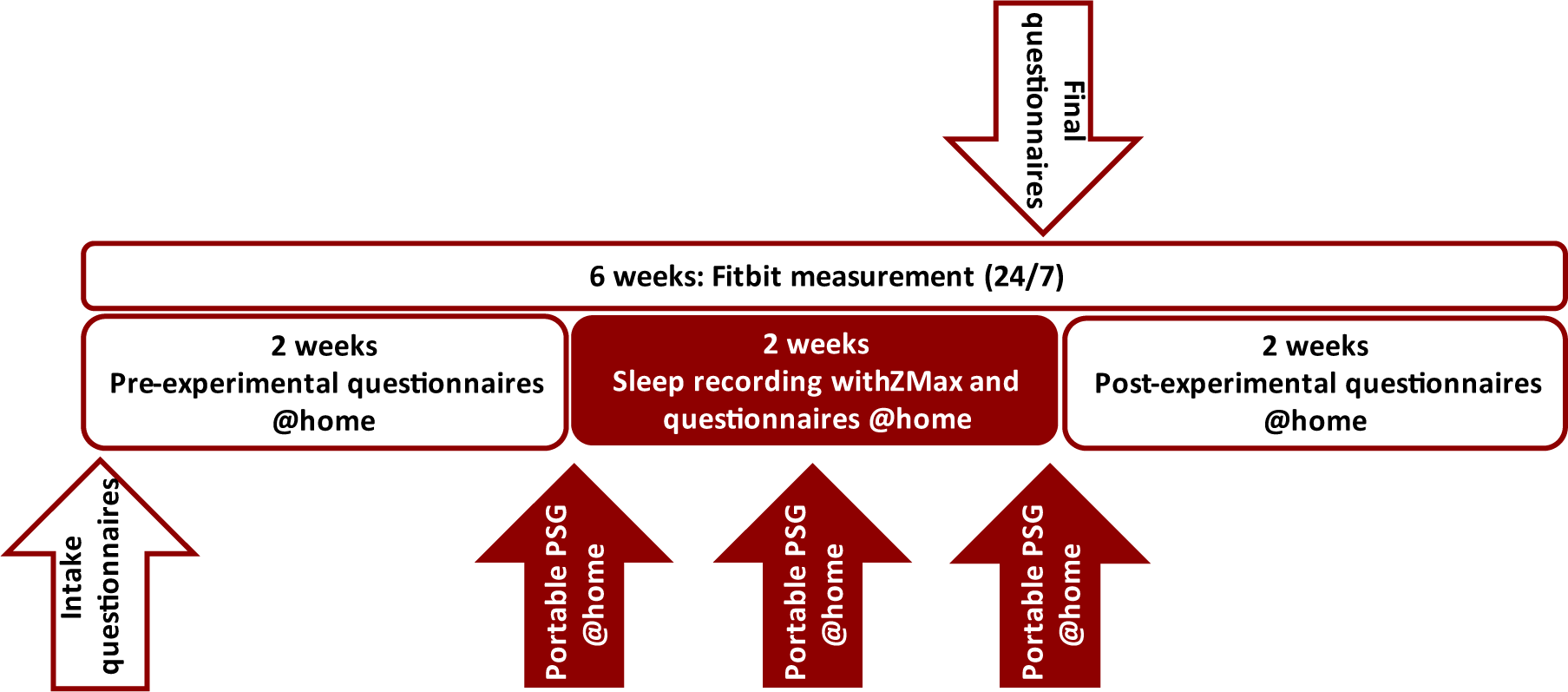
Longitudinal ZMax validation study design (dataset 2). This study was essentially designed for the validation of ZMax EEG headband concerning the PSG. Nocturnal sleep data were recorded during the middle two weeks (15 nights) using ZMax. There was a parallel PSG recording on the first, middle, and the last night of the middle two weeks.

As an indication of the lights-out time, the participants were instructed to follow a set of predefined tasks such as closing their eyes for one minute to relax, performing a series of teeth clenches, left-right-left-right (LRLR) eye movements, and several consecutive blinks. These events were also used to synchronize the data epochs recorded by the PSG and ZMax. In instances where a subject failed to perform the predefined tasks, alternative events (e.g., slow eye rollings during N1 sleep or movement arousals) were used for synchronization purposes.

### 2.4. Behavioral analysis

The behavioral analyses were conducted based on dataset 2 in which daily questionnaires were administered during pre-experimental, experimental, and post-experimental periods. Our analysis examined the influence of wearing the ZMax EEG headband on subjective sleep quality (on a scale from 0: very badly to 5: very well), number of awakenings, and mood (on a scale from 0: sad to 10: happy). In addition, the technology acceptance rate was evaluated through questions regarding the comfortability (on a scale from 0: very uncomfortable to 10: very comfortable) and sleep disturbance of the headband (on a scale from 0: not at all to 10: very much) during the experimental period.

We used JASP statistical software (Love et al., 2019) to conduct statistical analyses. A repeated measures ANOVA test with a post-hoc Holm correction was employed to detect significant differences between sleep quality, morning mood, and the number of awakenings during different study phases. For the remaining behavioral analyses, correlation values were calculated and plotted using JASP.

### 2.5. Preprocessing

Sleep scoring is the most fundamental step in performing any analysis of sleep data. Datasets 1-3 were scored by a single experienced professional, whereas dataset 4 was scored redundantly by a group of five university researchers with the most experienced scorer reviewing epochs with a lack of consensus and making a final decision on the basis of the 5 different scores and the respective data from those epochs. Datasets 1-3 were scored using Domino (SOMNOscreen ltd, Germany), whereas dataset 4 was scored using SpiSOP (https://www.spisop.org), all according to AASM protocols. Non-scorable PSG and their corresponding ZMax recordings were excluded from further analysis. A thorough examination of the simultaneous ZMax recordings was performed on the remaining scorable data. This process included the determination of artifacts that were present on the ZMax but not on the simultaneous PSG recording. During this process, the TFR of the ZMax EEG channels within the primary sleep frequency range of 0.3– 30 Hz was initially investigated to detect three types of issues: (1) ‘noisy epochs’, characterized by high power across a broad frequency range obscuring the identification of sleep microstructures such as 12– 15 Hz spindle or 0.5–4 Hz slow-oscillation (SO) activity, (2) ‘epochs of data loss’ indicated by the absence of power throughout the frequency range, and (3) ‘fragmented data’ containing several noisy or data loss epochs next to each other (see also Supplemental Figure 1). Subsequently, if the overall duration of artefactual and non-scorable epochs exceeded approximately 25% of a complete recording that consisted of a minimum 5.5 hours (as an estimate of a normal duration of nocturnal sleep), the data from that night were excluded from subsequent processing (see also Section 2.1). Otherwise, SpiSOP software was utilized to accurately mark the ‘noisy’ and ‘data loss’ epochs in the time domain. The remaining ‘clean’ epochs were retained for post-processing if a clear distinction between various sleep stages could be made, including high SO and spindle band power as an indication of N2 and slow-wave sleep (SWS), low spindle and mixed-frequency band power as an indication of restful wake, N1 or REM sleep, and brief bursts of high-power mixed-frequency activity as an indication of movement arousals (see also Supplemental Figure 1).

In case of different sampling rates between ZMax and standard PSG recordings, we resampled the data pairs to the one with a lower sampling frequency, i.e., 256 Hz in ZMax. To compare the analytic results derived from the data collected by ZMax and standard PSG, it was necessary to align the ground truth hypnograms (human scoring of the PSG data) with the corresponding ZMax recordings. This alignment was achieved by identifying a mutual event present in both sets of data, such as predefined eye movements. Depending on which device was started earlier, some epochs were either removed from the beginning of the hypnogram or added to the end (marked as non-scorable). The length of the hypnogram was then adjusted to match the number of epochs in ZMax recording, by either removing additional epochs or adding fake non-scorable epochs as needed.

### 2.6. Autoscoring

We compared the performance of two autoscoring approaches with the human scoring as the ground truth. The autoscoring algorithms encompass: (1) *ZLab,* which is the autoscoring service provided by ZMax Hypnodyne, and (2) the *DreamentoScorer* (Figure 3) from our open-source dream engineering toolbox (Esfahani et al., 2022a; https://github.com/dreamento/dreamento). To receive the results from ZLab, all the available ZMax data from datasets 1-4 were pseudonymized and then shared with Hypnodyne for autoscoring. While the proprietary architecture of ZLab remained undisclosed to us, it reportedly integrates various sources of information obtained by ZMax, including EEG, tri-axial accelerometer, and PPG data.

**Figure 3.**
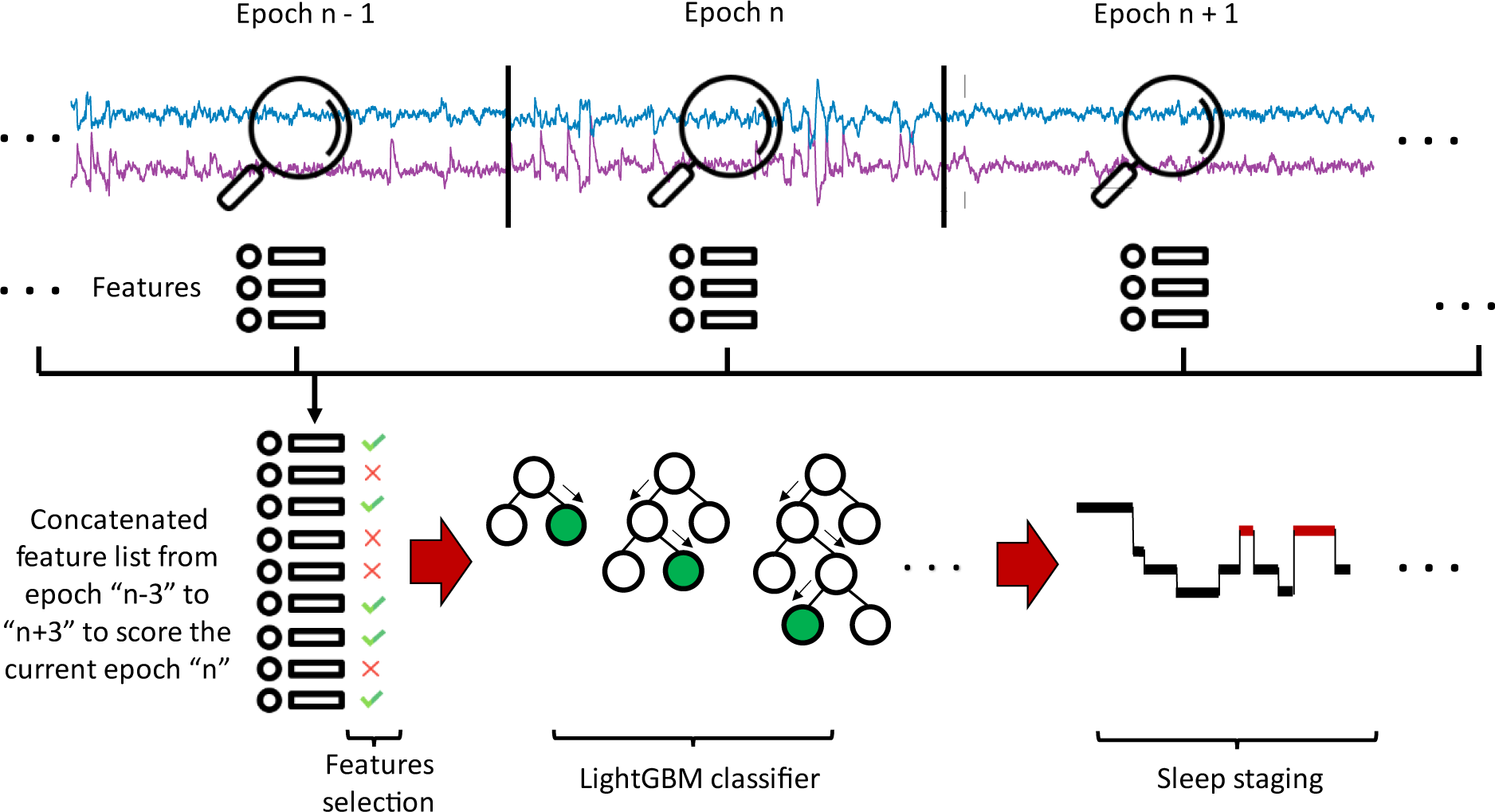
DreamentoScorer schematic representation. DreamentoScorer takes the temporal dependence between epochs into consideration. This means that, to score the current epoch, it extracts the features not only from the current epoch but also from 3 epochs back and forth. This allows for more accurate autoscoring using a simpler machine-learning model.

DreamentoScorer operates based on a LightGBM (Ke et al., 2017) machine learning model to automatically predict sleep stages. Initially, DreamentoScorer extracts a variety of time, frequency, and time-frequency domain features from each 30-second epoch data of both ZMax EEG channels (a full list of features can be found on Dreamento’s GitHub page). In light of importance of the temporal dependence for sleep scoring according to AASM protocols, DreamentoScorer extracts features from three epochs back and forth to provide an autoscoring result of the current epoch (see Figure 3). The Boruta algorithm (Kursa & Rudnicki, 2010) is then used to select the most relevant features to be used as inputs to the LightGBM classifier. The optimal list of DreamentoScorer hyperparameters were found through a random grid search (see Supplemental Table 1) and can be found on Dreamento GitHub page: https://github.com/dreamento/dreamento.

In this study, we applied a 10-fold cross-validation on datasets 2-4 (for dataset 1, due to the limited number of data we had to use 5-fold cross-validation), separately, and then across the pooled dataset. To assess the performance of autoscoring on dataset 5, due to the absence of PSG and thus human scoring, we visually investigated all the resulting hypnograms by DreamentoScorer and evaluated the alignment of the corresponding TFR events with the output scoring (see also the criteria for ‘clean’ data in section 2.5. Preprocessing as well as Supplemental Figure 1.A).

### 2.7. Sleep assessment metrics

Objective sleep assessment metrics proposed by AASM, including sleep period time (SPT), total sleep time (TST), sleep efficiency (SE), sleep maintenance efficiency (SME), sleep onset latency (SOL), wake after sleep onset (WASO), and the duration, percentage and latency of each sleep stage were derived based on the human scoring of the PSG to serve as the ground truth. These measures were then compared with the results obtained from the ZLab and DreamentoScorer autoscoring algorithms.

### 2.8. Bandpower analysis

We compared the power of the signals obtained from ZMax and PSG using the YASA toolbox (Vallat et al., 2021) across different frequency ranges, including SO (0.5–1 Hz), delta (1–4 Hz), theta (4–8 Hz), alpha (8– 12 Hz), and sigma (13–16 Hz) within different stages of sleep. The power of the signal was computed and averaged over F7-Fpz and F8-Fpz channels for ZMax and over EEG channels of the PSG in the proximity of ZMax i.e., F3:A2 and F4:A1 in datasets 1,2, 4, and F3 only (due to absence of F4) in dataset 3.

To further investigate the concordance of the power estimates between the measurement modalities, we employed Bland-Altman plots (Bland & Altman, 1986). Bland-Altman plots are suited to indicate the presence of systematic error (fixed bias) between the measurements as well as highlighting the outliers. Correlation analysis represents the degree to which two measurements are ‘associated’ (not the difference among measurements), whereas Bland-Altman plots demonstrate the level of ‘agreement’ between the measures (Giavarina et al., 2015).

### 2.9. Sleep microstructural analysis

Non-REM sleep microstructural features such as SOs and spindles and REM sleep microstructural features such as rapid eye movements were automatically detected in both ZMax and PSG signals using the algorithm proposed by Mölle et al. (2002) and Ngo et al., (2015), implemented in the SleepTrip toolbox (RRID: SCR_017318, https://github.com/Frederik-D-Weber/sleeptrip). For non-REM features, the events were detected using the closest channels of the PSG in the proximity of ZMax, namely F3:A2 and F4:A1 channels. To detect REM features, a derivation of the dedicated EOG channels of the PSG, i.e., horizontal EOG left - EOG right was compared with ZMax F7 - F8. The results were then averaged across channels for each system. The specific configuration and variables used for detecting these events can be found in the Supplemental Table 2. Raincloud plots (Allen et al., 2019) were employed to demonstrate the agreement between the measures from ZMax vs. PSG signal.

We have also compared the morphology of the detected events in ZMax vs PSG by illustrating the event-related potential (ERP). To create the ERP of the SOs and spindles, the detected events were time-locked to the maximum absolute trough and then averaged across trials. The characteristics of microstructural features such as counts, density, duration, and amplitude were then analyzed and the correlation between ZMax and the PSG outputs was determined. Additionally, Bland-Altman plots were employed to depict the measurement agreement between the PSG and ZMax.

## 3. Results

The behavioral analyses were performed on 441, 462, and 403 nights during pre-experimental, experimental, and post-experimental phases, respectively, which accounted for an average of 13.78 ± 0.97, 14.44 ± 1.08, 12.59 ± 1.04 nights per participant. We found no striking difference in subjective sleep quality when comparing the pre-experimental and experimental phases (t = 1.80, p = 0.076), whereas sleep quality improved during the post-experimental phase, both in comparison with the experimental (d = -0.51, p < 0.001) and pre-experimental phase (d = -0.29, p = 0.04) (Figure 4-A, Supplemental Table 3). Nevertheless, the average sleep quality during all phases remained between the ‘fairly well’ and ‘well’ grades (3.29 ± 0.66, 3.15 ± 0.66, 3.48 ± 0.64 for pre-experimental, experimental, and post-experimental phases, respectively, where 3 represents ‘fairly well’ and 4 is ‘well’). Furthermore, morning mood did not differ across the different phases of the study between pre-, post-, and experimental phases (ppre- vs. exp. = 0.86, ppre- vs. post-exp = 0.19, ppost- vs. exp= 0.19) as demonstrated in Figure 4-B and Supplemental Table 3. In the post-experimental period the number of awakenings were reduced compared to both the pre-experimental (d = 0.44, p = 0.005) and experimental phases (d = 0.54, p < 0.001). Although this difference is statistically significant, there was only a slight difference in average number of awakenings observed: 1.39 ± 1.00, 1.49 ± 0.05, and 0.98 ± 0.83, during the pre-experimental, experimental, and post-experimental phases, respectively (Figure 4-C). Focusing on the experimental period in particular, regressional trends were mainly absent for sleep quality (r = 0.09, p = 0.053, Figure 4-D), morning mood (r = 0.02, p = 0.724, Figure 4-E), number of awakenings (r = -0.06, p = 0.223, Figure 4-F), ZMax disturbance (r = -0.03, p = 0.54, Figure 4-G), and ZMax comfort (r = 0.05, p = 0.32, Figure 4-H) over the course of 15-night experimental period.

**Figure 4.**
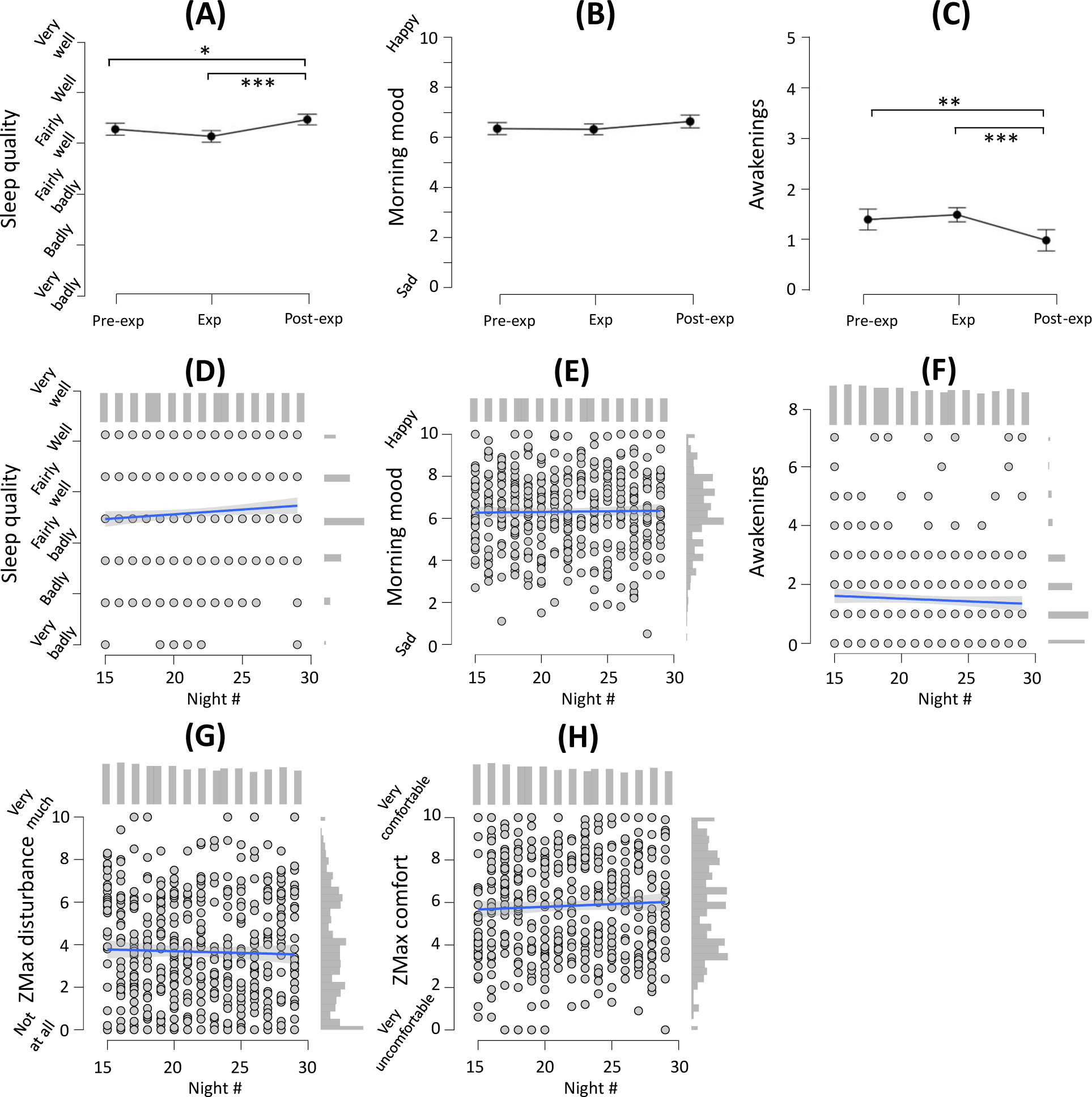
Results of behavioral analysis based on dataset 2. **(A)** subjective sleep quality, **(B)** morning mood, and **(C)** number of awakenings across different phases of the study. Panels **(D)** - **(H)**, focus on the experimental period only, i.e., from nights 15 - 29. **(D)** subjective sleep quality, **(E)** morning mood (n.s.), **(F)** the number of awakenings (n.s.), **(G)** ZMax disturbance (n.s.), and **(H)** ZMax comfort (n.s.) trends during the experimental period. The bars on the right of x- and top of y-axes of panels (D) - (H) represent the histograms. n.s. : not significant, pre-exp: pre-experimental phase, exp: experimental phase, post-exp: post-experimental phase. ***: p-value < .001, **: p-value < .01.

To demonstrate the similarity of the recorded signals by ZMax and the PSG, a sample 30-second epoch from each sleep stage is illustrated in Figure 5. During wakefulness, signals recorded during restful wake with eyes closed, while performing predefined eye signaling, and during the events of jaw clenching exhibited relatively similar patterns. This is important as these signals could be used to synchronize ZMax output signals with simultaneous recording from another measurement modality. For instance, predefined eye movements are useful for synchronization with another EEG system, whereas jaw clenching events might be employed to synchronize with a simultaneous EMG recording. During N1 sleep, particularly during a period of slow eye-rolling, ZMax signal displays a combination of the EOG (manifested as the slow/low-frequency baseline activity) in addition to the EEG (evidenced by the low-amplitude mixed-frequency activity). Furthermore, microstructural features characterizing non-REM sleep, such as sleep spindles and K-Complexes during N2 sleep as well as SOs during SWS and rapid eye movement events during REM sleep are also discernible within the ZMax signal.

**Figure 5.**
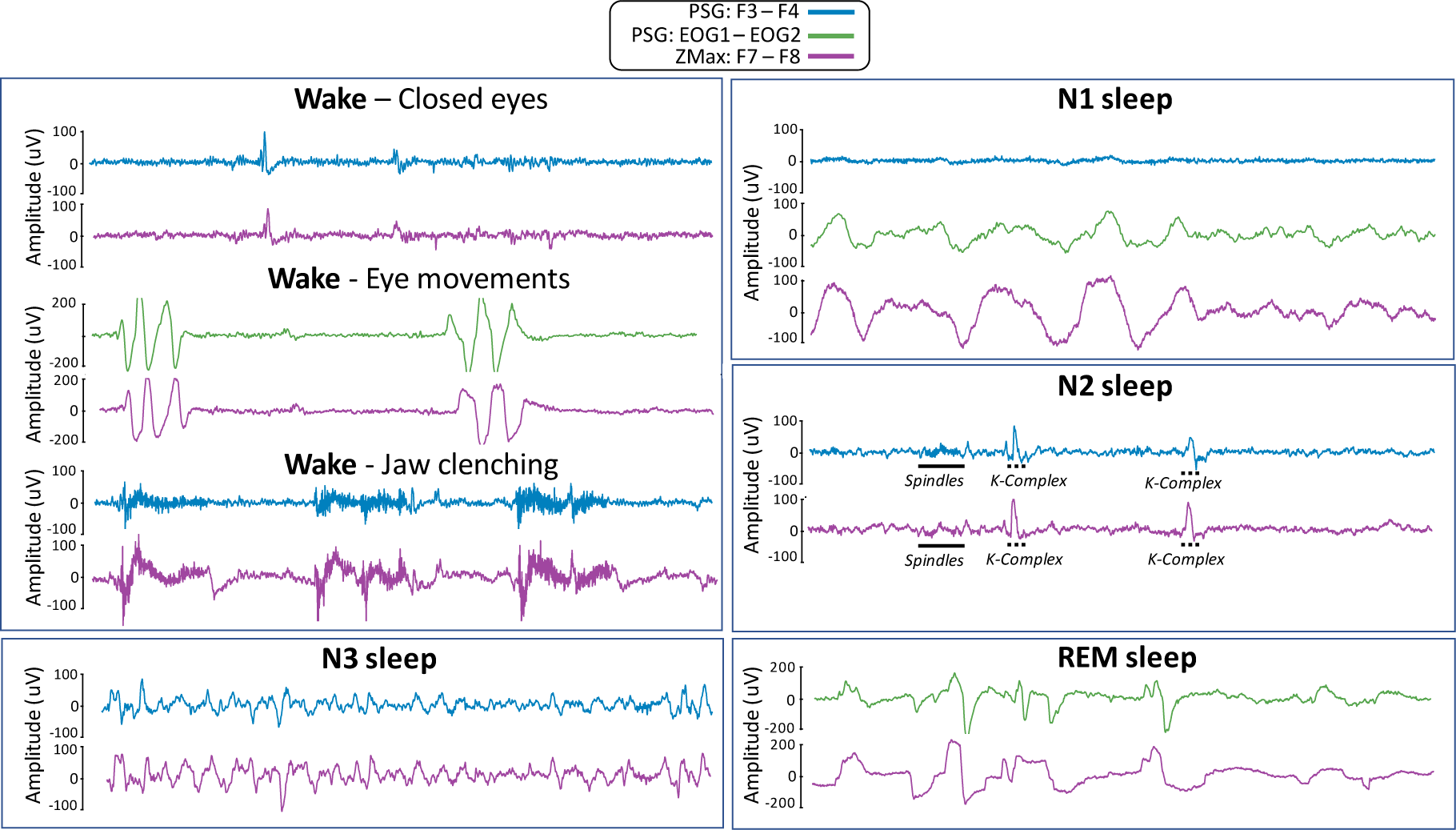
Sample representation of a simultaneous PSG and ZMax recording during different stages of sleep from a single participant. During wakefulness, comparison between the ZMax and PSG signals were made while (1) resting with the eyes closed, (2) performing pre-defined left-right-left-right (LRLR) eye signaling as an event to synchronize signals between the PSG and ZMax, and (3) performing jaw clenching as an alternative event for synchronization between the measurement modalities. The ZMax vs. PSG signals were also compared within N1 sleep, N2 sleep, N3**/**SWS, and REM sleep. The F3 - F4 derivation from the PSG (blue), was compared with the F7 - F8 derivation from ZMax (purple). An additional EOG derivation, i.e., EOG left - EOG right (green) was also depicted to compare slow eye rolling during N1 and rapid eye movement during REM sleep. All illustrations show a 30-s epoch of data. The y-xes show m litu es in μV.

In an early stage of this study, when we only had datasets 2 and 3 manually scored by our human experts, we tested the performance of one of the well-known algorithms on our wearable system data. We applied U-sleep v1.0 (Perslev et al., 2021) as one of the most comprehensive autoscoring algorithms including flexibility in terms of the chosen EEG montage, to 69 pre-processed ZMax recordings. Our results indicated that if U-sleep is employed ‘out of the box’, without any fine tuning (e.g., training partially based on wearable data) and using a double EEG input in place of the typical EEG+EOG it would result in an overall accuracy = 6 .3%, mF1 = 0.63%, and a Cohen’s kappa score = 57.3%, which is lower than both the human inter-rater agreements as well as its performance on PSG data. As expected, fine-tuning through applying a 10-fold cross-validation (similarly, using double EEG inputs) improved the results, achieving overall accuracy = 7 .6%, mF1 = 72.7%, Cohen’s kappa score = 71%.

We also evaluated the performance of Zlab and DreamentoScorer as wearable-specific autoscoring algorithms for our four datasets, including 135 data comprising over 900 hours of night sleep (108064 epochs, after artifactual epochs rejection). The performance of the algorithms for sleep autoscoring was analyzed and compared across various datasets, as detailed in Table 2. We established the feasibility of developing an autoscoring model (DreamentoScorer) based on ZMax data which may also be utilized by future ZMax studies. When compared to the ground truth scoring, DreamentoScorer consistently outperforms Zlab across all datasets. More specifically, DrementoScorer achieved a Cohen’s kappa score of 64.84%, 71.60%, 64.39%, 83.00%, and 72.18% for datasets 1, 2, 3, 4, and the pooled dataset, surpassing the Cohen’s kappa scores of 59.40%, 57. 9%, 61.4 %, 61.34%, and 59.61% obtained from ZLab. Furthermore, when delving into the sleep stage classification of the pooled dataset, Dreamento yielded higher F1-scores compared to ZLab. Specifically, for wake, N1, N2, SWS, and REM sleep, Dreamento achieved F1-scores of 77.36%, 39.73%, 81.23%, 82.28%, 83.01%, whereas ZLab obtained 74.67%, 24.67%, 73.59%, 69.05%, and 76.30%, respectively.

**Table 2.** Comparison between the ZLab and DreamentoScorer autoscoring algorithms concerning the human scoring results as the ground truth. Recall = tp / (tp + fn), Precision = tp / (tp + fp), F1-score = 2 * (precision * recall) / (precision + recall), Accuracy =(tp + tn) / (tp + fp + tn + fn), macro-F1 (mF1) = the mean of F1-score, ohen’s k score κ = Po - Pe) / (1 - Pe) where: Po is the observed, and Pe is the expected agreement. The values are reported as percentages. The bolded values show the higher outcome derived from either ZLab or DreamentoScorer.

|  | Dataset | Wake |  | N1 |  | N2 |  | SWS |  | REM |  |
| --- | --- | --- | --- | --- | --- | --- | --- | --- | --- | --- | --- |
|  |  | ZLab | Dreamento | ZLab | Dreamento | ZLab | Dreamento | ZLab | Dreamento | ZLab | Dreamento |
| Recall (%) | Dataset 1<br>(N =9) | 70.39 | **80.33** | 26.31 | **44.63** | 76.63 | **81.22** | **80.90** | 64.83 | 77.45 | **84.38** |
| Precision (%) |  | 41.78 | **61.44** | 45.30 | **51.57** | 84.10 | **84.18** | 56.95 | **67.24** | **79.89** | 77.71 |
| F1-score (%) |  | 52.44 | **69.63** | 33.29 | **47.85** | 80.19 | **82.67** | 66.84 | **66.01** | 78.65 | **80.91** |
| Overall Accuracy (%) |  | **ZLab: 71.10 , Dreamento: 75.27** |  |  |  |  |  |  |  |  |  |
| Overall mf1(%) |  | **ZLab: 62.28, Dreamento: 69.41** |  |  |  |  |  |  |  |  |  |
| Overall Kappa (%) |  | **ZLab: 59.40, Dreamento: 64.84** |  |  |  |  |  |  |  |  |  |
| Recall (%) | Dataset 2<br>(N = 66) | 75.22 | **78.45** | 29.12 | **33.73** | 75.16 | **75.33** | 59.42 | **83.63** | 75.14 | **85.60** |
| Precision (%) |  | **79.73** | 75.07 | 14.69 | **39.37** | 63.72 | **76.98** | 86.23 | **88.20** | 68.18 | **76.29** |
| F1-score (%) |  | **77.41** | 76.72 | 19.53 | **36.33** | 68.97 | **76.15** | 70.36 | **85.86** | 71.49 | **80.68** |
| Overall Accuracy (%) |  | **ZLab: 68.21 , Dreamento: 78.87** |  |  |  |  |  |  |  |  |  |
| Overall mf1(%) |  | **ZLab: 61.55 , Dreamento: 71.15** |  |  |  |  |  |  |  |  |  |
| Overall Kappa (%) |  | **ZLab: 57.89, Dreamento: 71.60** |  |  |  |  |  |  |  |  |  |
| Recall (%) | Dataset 3<br>(N = 17) | 64.61 | **76.58** | 16.92 | **22.05** | **82.09** | 72.93 | 73.66 | **83.73** | **73.90** | 73.10 |
| Precision (%) |  | **85.61** | 70.49 | 22.76 | **35.53** | 67.63 | **78.10** | **80.71** | 76.43 | **70.17** | 69.98 |
| F1-score (%) |  | **73.64** | 73.40 | 19.41 | **27.21** | 74.16 | **75.43** | 77.03 | **79.91** | **71.99** | 71.50 |
| Overall Accuracy (%) |  | **ZLab: 71.46 , Dreamento: 73.18** |  |  |  |  |  |  |  |  |  |
| Overall mf1(%) |  | **ZLab: 63.25 , Dreamento: 65.49** |  |  |  |  |  |  |  |  |  |
| Overall Kappa (%) |  | **ZLab: 61.48 , Dreamento: 64.39** |  |  |  |  |  |  |  |  |  |
| Recall (%) | Dataset 4<br>(N = 43) | 84.40 | **86.87** | 41.48 | **57.49** | 66.75 | **89.65** | **92.79** | 90.02 | 82.34 | **94.02** |
| Precision (%) |  | 61.68 | **85.76** | 33.72 | **64.07** | 91.56 | **94.28** | 47.41 | **81.44** | **89.12** | 87.93 |
| F1-score (%) |  | 71.42 | **86.31** | 37.20 | **60.61** | 77.21 | **91.91** | 62.75 | **85.52** | 85.60 | **90.87** |
| Overall Accuracy (%) |  | **ZLab: 73.90, Dreamento: 89.49** |  |  |  |  |  |  |  |  |  |
| Overall mf1(%) |  | **ZLab: 66.83, Dreamento: 83.04** |  |  |  |  |  |  |  |  |  |
| Overall Kappa (%) |  | **ZLab: 61.34, Dreamento: 83.00** |  |  |  |  |  |  |  |  |  |
| Recall (%) | Pooled data<br>(N = 135) | 73.73 | **79.13** | 28.86 | **37.10** | 72.86 | **78.33** | 67.74 | **83.54** | 77.28 | **87.65** |
| Precision (%) |  | 75.63 | **75.66** | 21.54 | **42.77** | 74.32 | **84.35** | 70.41 | **81.06** | 75.35 | **78.84** |
| F1-score (%) |  | 74.67 | **77.36** | 24.67 | **39.73** | 73.59 | **81.23** | 69.05 | **82.28** | 76.30 | **83.01** |
| Overall Accuracy (%) |  | **ZLab: 70.50 , Dreamento: 79.67** |  |  |  |  |  |  |  |  |  |
| Overall mf1(%) |  | **ZLab: 63.65 , Dreamento: 72.72** |  |  |  |  |  |  |  |  |  |
| Overall Kappa (%) |  | **ZLab: 59.61 , Dreamento: 72.18** |  |  |  |  |  |  |  |  |  |

Additionally, the ground truth hypnogram (Figure 6-A, top) was schematically compared for a sample data with the hypnograms resulting from ZLab (Figure 6-A, middle) and DreamentoScorer (Figure 6-A, bottom). To gauge more details about the performance of DreamentoScorer in classifying each 30-second epoch of data, the classifier’s probability diagram (hypodensity) is shown in Figure 6–B (hypodensity was not provided for ZLab autoscoring) and the TFRs from both ZMax EEG channels were demonstrated in Figure 6-C.

**Figure 6.**
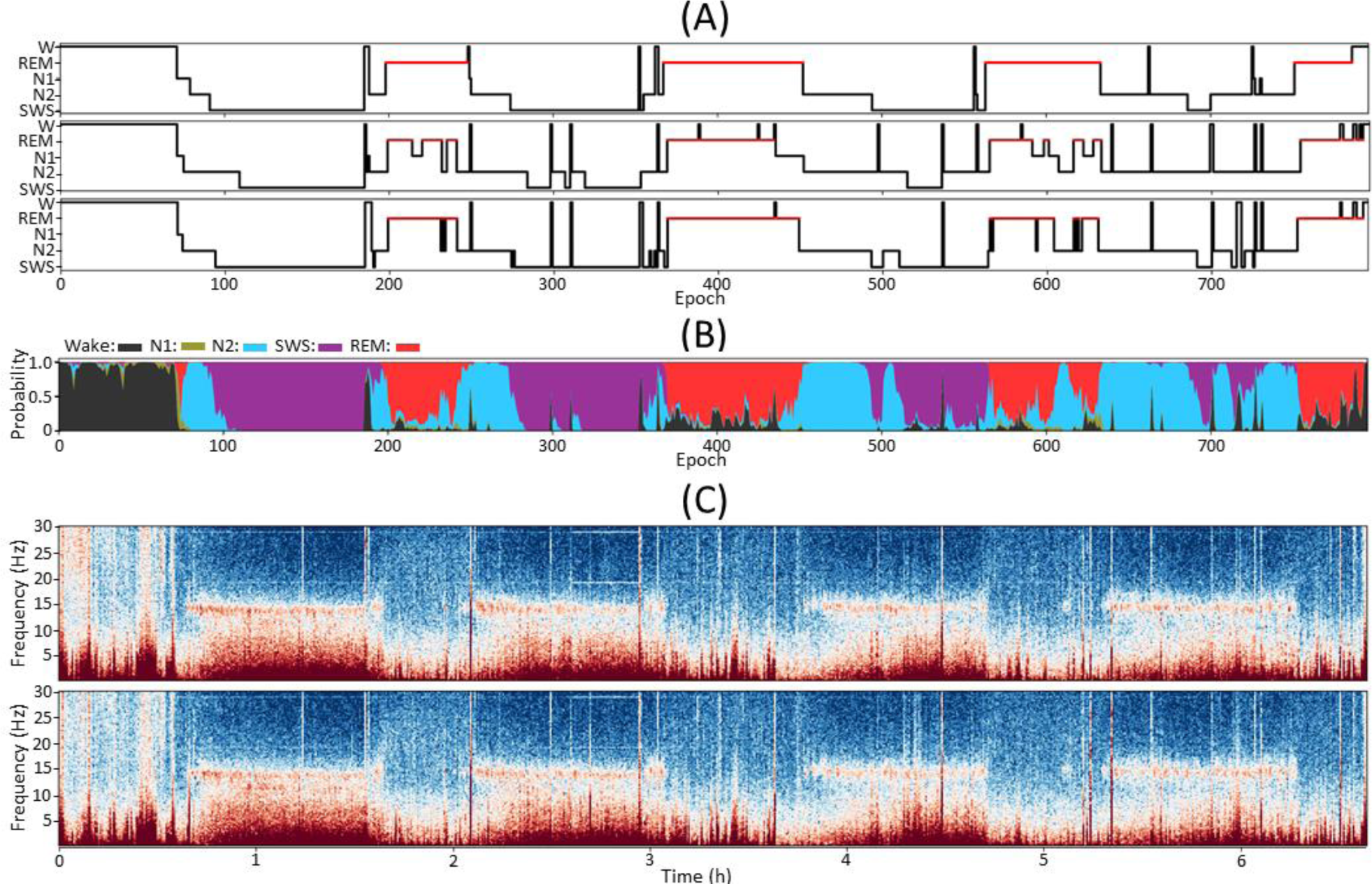
Comparison between the performance of autoscoring algorithms with the ground truth based on human scoring of the PSG. **(A)** ground truth hypnogram (top), ZLab hypnogram (middle), and Dreamento hypnogram (bottom). **(B)** Hypnodensity diagram of Dreamento representing probability of each sleep stage while predicting each epoch of data by DreamentoScorer, with different colors representing different stages of sleep, black: Wake, green: N1, light blue: N2, purple: SWS, red: REM. **(C)** the time-frequency representation of the F7-Fpz, and F8-Fpz channels from ZMax.

Datasets 1-4 were used in this study to benchmark ZMax performance across healthy young adults, whereas dataset 5 was included merely as a proof of concept to show the potential applicability of ZMax and DreamentoScorer over (1) an older population (2) with pathological conditions (see Table 1). While the detailed analysis for dataset 5 will be discussed elsewhere (ter Avest et al., paper in preparation), here we report that according to our data quality assessment protocols (Supplemental Figure 1), out of 512 available nocturnal sleep recordings from IBD patients, 280 (55%) were deemed entirely useful, 366 (71%) contained at least 75% useful data, 83 (17%) included fragmented data, and 63 (12%) only contained noise. Given the absence of ground truth PSG to assess autoscoring for dataset 5, we defined a shallow analysis as follows: we first autoscored the entirely useful portion of data and then visually inspected whether the autoscoring results are aligned with the sleep stage transition criteria through TFR (see Figure 6 and Supplemental Figure 1). Based on this shallow analysis, we observed that autoscoring using DreamentoScorer resulted in acceptable outcomes in 82% of cases. An explanation for the remaining poor autoscorings, could be the subjects’ age range (46. 7 ± 13.15 years of age for acceptable autoscoring results vs 54.19 ± 11.96 years of age for poor autoscoring results), underlying disease and potential comorbidities.

Based on the manual scoring and autoscoring results, we calculated various sleep assessment metrics suggested by AASM. The results derived from the manual scoring served as the ground truth, and were compared with the outcomes from the autoscoring approaches (see Table 3). Overall, we observed acceptable outcomes for both ZLab and DreamentoScorer, when compared to manual scoring (see also the raincloud plot and statistical assessment in Supplemental Figure 2).

**Table 3.** Comparative sleep assessment metrics. SPT: sleep period time, TST: total sleep time, SE: sleep efficiency, SME: Sleep maintenance efficiency, SOL: sleep onset latency, WASO: wake after sleep onset, Lat: latency. Values are represented in the form of mean ± standard deviation.

| Dataset |  |  |  |  |  |  |  |  |  |  |  |  |  |  |  |
| --- | --- | --- | --- | --- | --- | --- | --- | --- | --- | --- | --- | --- | --- | --- | --- |
|  | Dataset 1: Donders 2018 |  |  | Dataset 2: Donders 2022 |  |  | Dataset 3: Stockholm University |  |  | Dataset 4: QS |  |  | Pooled data |  |  |
|  | Ground truth | ZLab | Dreamento | Ground truth | ZLab | Dreamento | Ground truth | ZLab | Dreamento | Ground truth | ZLab | Dreamento | Ground truth | ZLab | Dreamento |
| **SPT (min)** | 431.5±72.03 | 422.94±85.09 | 435.22±69.09 | 392.09±71.5 | 390.42±70.69 | 395.14±72.78 | 365.65±66.1 | 373.79±70.06 | 368.18±65.37 | 348.52±34.39 | 348.66±34.3 | 348.67±34.34 | 377.49±65.95 | 377.18±66.32 | 380.66±68.74 |
| **TST (min)** | 415.28±73.74 | 397.56±76.34 | 411.11±72.64 | 363.56±65.13 | 366.86±65.24 | 360.94±65.55 | 331.15±64.48 | 349.35±67.09 | 324.74±61.6 | 341.95±33.46 | 336.91±31.63 | 341.78±34.12 | 356.04±61.00 | 357.15±60.37 | 354.30±61.37 |
| **SE (%)** | 93.96±4.31 | 89.74±6.58 | 92.03±6.8 | 86.5±6.38 | 87.23±5.67 | 85.88±6.67 | 81.74±6.27 | 86.27±6.52 | 80.48±9.18 | 96.27±3.23 | 94.86±2.69 | 96.2±3.39 | 89.51±7.57 | 89.71±6.26 | 89.02±8.22 |
| **SME (%)** | 96.12±3.41 | 94.2±1.57 | 94.37±5.89 | 93.0±5.63 | 94.06±3.29 | 91.58±5.53 | 90.52±5.23 | 93.48±3.04 | 88.33±7.27 | 98.18±3.08 | 96.71±2.45 | 98.07±3.24 | 94.55±5.51 | 94.84±3.19 | 93.39±6.39 |
| **SOL (min)** | 8.22±5.95 | 17.67±24.68 | 10.22±6.57 | 25.7±15.94 | 30.6±24.11 | 23.67±19.5 | 25.82±15.97 | 28.15±22.11 | 23.88±17.95 | 5.64±2.19 | 5.72±2.66 | 5.63±1.93 | 18.16±15.87 | 21.5±22.69 | 15.73±15.75 |
| **WASO (min)** | 16.22±13.36 | 25.39±10.66 | 24.11±25.32 | 28.53±27.01 | 23.57±15.31 | 34.2±26.88 | 34.5±20.39 | 24.44±11.79 | 43.44±30.19 | 6.57±11.59 | 11.76±9.35 | 6.9±11.76 | 21.45±24.11 | 20.03±14.11 | 26.36±27.98 |
| **N1 (min)** | 51.94±26.78 | 30.17±12.04 | 46.11±27.59 | 15.92±12.56 | 31.55±21.84 | 13.64±13.12 | 25.21±14.32 | 18.74±13.1 | 15.65±8.32 | 11.33±6.46 | 13.93±7.38 | 10.16±7.2 | 18.09±16.28 | 24.23±18.65 | 15.72±12.01 |
| **N1 (%)** | 12.21±5.62 | 7.52±2.45 | 10.88±5.73 | 4.34±3.41 | 8.46±5.27 | 3.75±3.51 | 7.68±4.34 | 5.41±3.86 | 4.74±2.6 | 3.28±1.83 | 4.1±2.08 | 2.98±2.06 | 4.96±4.10 | 6.62±4.60 | 4.37±3.07 |
| **N2 (min)** | 208.22±48.99 | 189.72±34.38 | 201.22±53.56 | 128.58±42.93 | 151.66±55.62 | 125.83±41.55 | 141.56±36.39 | 171.82±52.7 | 132.18±50.82 | 201.76±27.71 | 147.08±51.5 | 191.84±28.69 | 158.38±51.08 | 155.27±54.10 | 147.53±51.05 |
| **N2 (%)** | 50.32±10.8 | 48.68±8.54 | 48.85±10.61 | 35.1±8.58 | 41.54±13.57 | 34.58±8.75 | 42.56±5.69 | 48.93±11.36 | 40.14±11.88 | 58.98±5.13 | 43.75±14.67 | 56.06±5.12 | 44.57±12.94 | 43.65±13.68 | 41.66±13.34 |
| **SWS (min)** | 60.5±19.03 | 85.94±36.09 | 58.33±31.73 | 139.92±33.64 | 96.42±46.52 | 132.67±37.07 | 101.94±24.92 | 93.03±35.92 | 111.68±43.99 | 54.99±11.4 | 107.63±48.93 | 60.78±13.61 | 102.76±47.09 | 98.87±45.96 | 105.94±47.25 |
| **SWS (%)** | 15.39±6.62 | 21.4±7.38 | 15.35±10.89 | 38.95±9.25 | 26.82±13.41 | 37.34±10.9 | 31.06±5.83 | 27.26±10.47 | 35.57±15.45 | 16.1±3.07 | 31.87±13.75 | 17.91±4.07 | 29.10±12.94 | 28.12±13.19 | 30.35±14.10 |
| **REM (min)** | 94.61±36.69 | 91.72±38.87 | 105.44±40.71 | 79.14±26.35 | 87.22±44.58 | 88.8±26.99 | 62.44±22.08 | 65.76±22.09 | 65.24±29.38 | 73.88±18.90 | 68.27±16.76 | 79.0±18.47 | 76.82±25.74 | 78.78±36.41 | 85.12±29.99 |
| **REM (%)** | 22.08±6.21 | 22.4±6.63 | 24.91±6.69 | 21.6±5.63 | 23.19±9.65 | 24.33±5.43 | 18.7±4.78 | 18.4±4.03 | 19.55±7.47 | 21.64±5.08 | 20.28±4.58 | 23.05±4.46 | 21.37±5.41 | 21.6±7.77 | 23.62±6.18 |
| **N1 Lat (min)** | 8.61±5.68 | 17.67±24.68 | 10.22±6.57 | 27.14±16.72 | 36.61±35.56 | 35.75±39.61 | 25.82±15.97 | 46.0±55.84 | 36.24±35.06 | 5.64±2.19 | 10.13±29.62 | 5.67±1.95 | 18.89±16.58 | 28.09±38.94 | 22.26±33.69 |
| **N2 lat (min)** | 23.61±25.5 | 23.72±25.64 | 15.83±4.87 | 33.79±23.38 | 35.88±25.59 | 35.24±25.05 | 31.35±19.42 | 31.94±20.66 | 33.35±19.78 | 8.7±2.64 | 8.3±2.58 | 9.21±2.65 | 24.81±22.09 | 25.79±23.92 | 24.59±20.93 |
| **SWS lat (min)** | 39.56±23.63 | 39.06±27.79 | 40.39±17.18 | 39.64±25.6 | 51.11±30.21 | 42.15±36.4 | 41.12±18.87 | 47.21±21.72 | 42.76±20.73 | 24.34±3.78 | 20.53±8.44 | 24.53±5.07 | 34.94±21.45 | 40.07±27.71 | 35.03±27.68 |
| **REM lat (min)** | 101.94±54.89 | 96.56±43.13 | 93.89±55.06 | 128.48±57.8 | 83.1±32.74 | 70.64±56.13 | 108.74±42.32 | 88.0±41.21 | 56.82±47.83 | 95.23±37.89 | 85.66±33.79 | 80.9±37.03 | 113.56±52.53 | 85.42±35.17 | 74.79±50.27 |

Our results from Table 3 as well as Supplemental Figure 2 demonstrate that the correlation between the ground truth and the results from ZLab and DreamentoScorer are rather close for measuring SPT (rZLab = 0.98 and r = 0.99, p-values < 0.001), TST (rZLab = 0.97 and rDreamento = 0.96, p-values < 0.001), SE (both r = 0.87, p-values < 0.001), SME (both r = 0.79, p-values < 0.001), SOL (rZLab = 0.86 and rDreamento = 0.81, p-values < 0.001), and WASO (rZLab = 0.79 and rDreamento = 0.81, p-values < 0.001). Compared with ZLab, DreamentoScorer resulted in higher correlation to the ground truth for determining the majority of stage-wise metrics, namely the duration of N1 (rZLab = 0.19, p-value < 0.05 rDreamento = 0.57, p-values < 0.001), N1 % (rZLab = 0.09 [n.s.], rDreamento = 0.51, p-values < 0.001), N2 (rZLab = 0.23, rDreamento = 0.79, p-values < 0.001), N2 % (rZLab = 0.09 [n.s.], rDreamento = 0.65, p-values < 0.001), SWS (rZLab = 0.06 [n.s.], rDreamento = 0.69, p-values < 0.001), SWS % (rZLab = 0.04 [n.s.], rDreamento = 0.63, p-values < 0.001), whereas REM duration (rZLab = 0.59, rDreamento = 0.71, p-values < 0.001) and REM % (rZLab = 0.38, rDreamento = 0.53, p-values < 0.001) resulted in more similar outputs between the algorithms. Additionally, ZLab and DreamentoScororer exhibited similar level of correlation to the ground truth when measuring N1 latency (rZLab = 0.49 and rDreamento = 0.41, p-values < 0.001), N2 latency (rZLab = 0.93 and rDreamento = 0.89, p-values < 0.001), and SWS latency (rZLab = 0.83 and rDreamento = 0.84, p-values < 0.001), nonetheless, the resulting REM latency (rZLab = 0.38, p-values < 0.001, rDreamento = 0.20, p-values < 0.05) better correlated with ZLab outcome.

By analyzing the agreement between ZMax and PSG in terms of bandpower computation using Bland-Altman plots, we found that ZMax exhibits a relatively low mean bias when compared to PSG. Overall, with respect to PSG relative bandpower, ZMax demonstrated the bias of -0.055 ± 0.085, 0.045 ± 0.044, 0.016 ± 0.027, 0.005 ± 0.012, -0.008 ± 0.023, +0.002 ± 0.002, and 1.13e-10 ± 2.22e-10 for delta, theta, alpha, sigma, beta, gamma, and absolute overall power, respectively (see also Supplemental Table 4 and 5). Nevertheless, the proportional bias of the ZMax concerning PSG for absolute power is clearly evident in Figure 7. Moreover, although a linear trend indicating a proportional bias was observed for higher frequencies such as the beta and gamma bands, this was limited to datasets 1 and 4, where the same headbands (earlier versions of ZMax) were utilized (see also Supplemental Figure 3).

**Figure 7.**
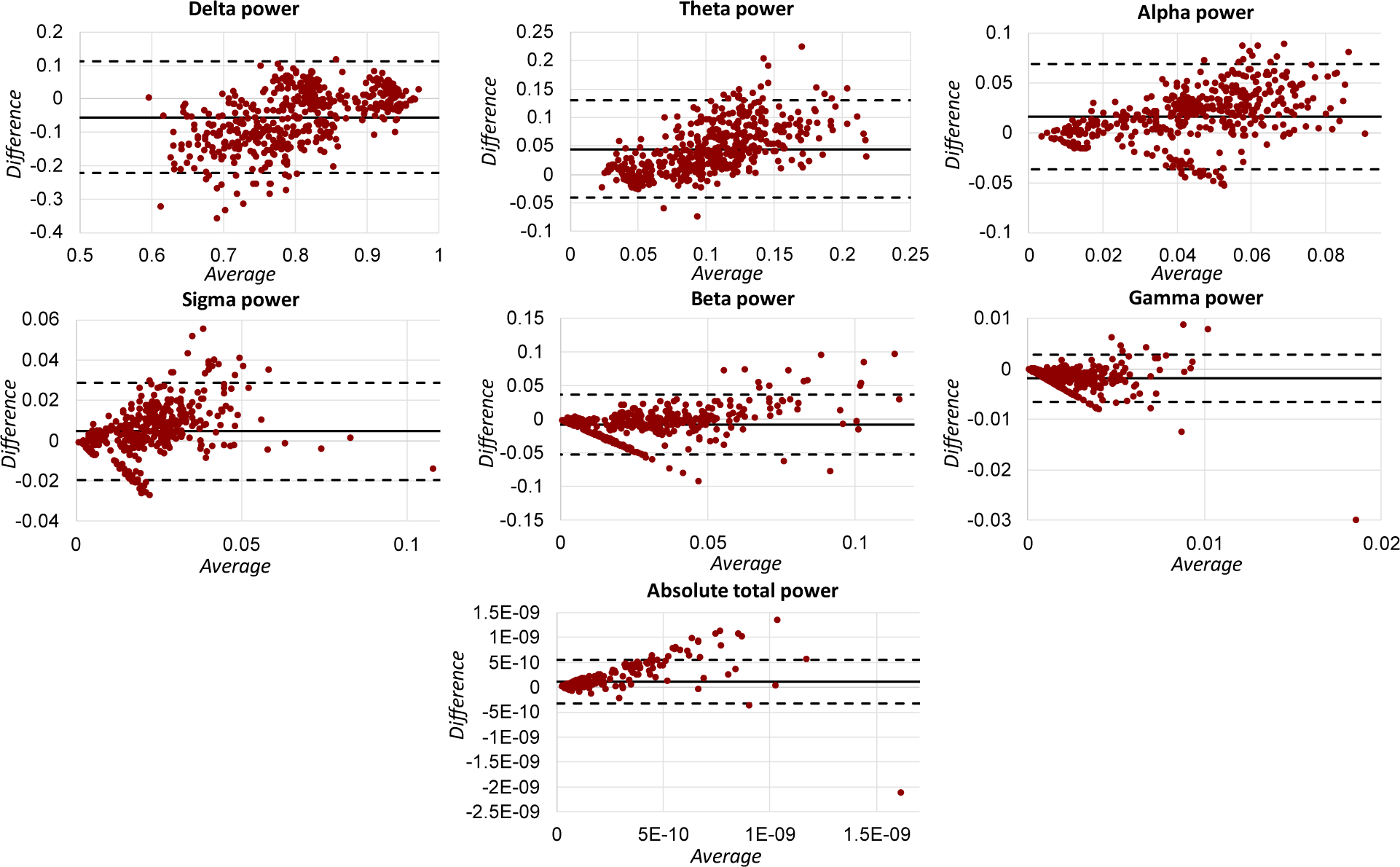
Bland-Altman plots representing the agreement between the relative bandpower between PSG and ZMax. The horizontal solid line in each figure represents the mean bias. The two dashed horizontal lines show the upper and lower limits of agreement. The results are demonstrated based on the pool dataset. The difference between the measures was based on PSG outcome minus the ZMax-based measure. In each plot, the x-axis represents the average of the two measures, whereas the difference between measures is shown on the y-axis.

Focusing on the number of detected events, ZMax consistently underestimated the number of detected SO and spindle counts compared to PSG (165.62 ± 92.00 vs. 404.43 ± 162.40, and 919.97 ± 357.53 vs. 1008.7 ± 255.11, for SO and spindle counts over the pooled dataset, respectively). This can also be schematically represented alongside the hypnogram as in Supplemental Figure 4. Conversely, ZMax overestimated the number of detected REM events (831.57 ± 549.31 vs. 467.01 ± 511.6 over the pooled dataset). We further observed an underestimation of SO density, SO frequency, SO trough-to-peak amplitude and slope, spindle density, spindle duration, spindle trough-to-peak amplitude, spindle linear regression frequency, and REM mean speed. Conversely, the results derived from ZMax overestimated SO mean duration, SO zero-crossing slope, REM counts, REM density, and REM duration (see Figure 8 and Supplemental Table 6 for more details).

**Figure 8.**
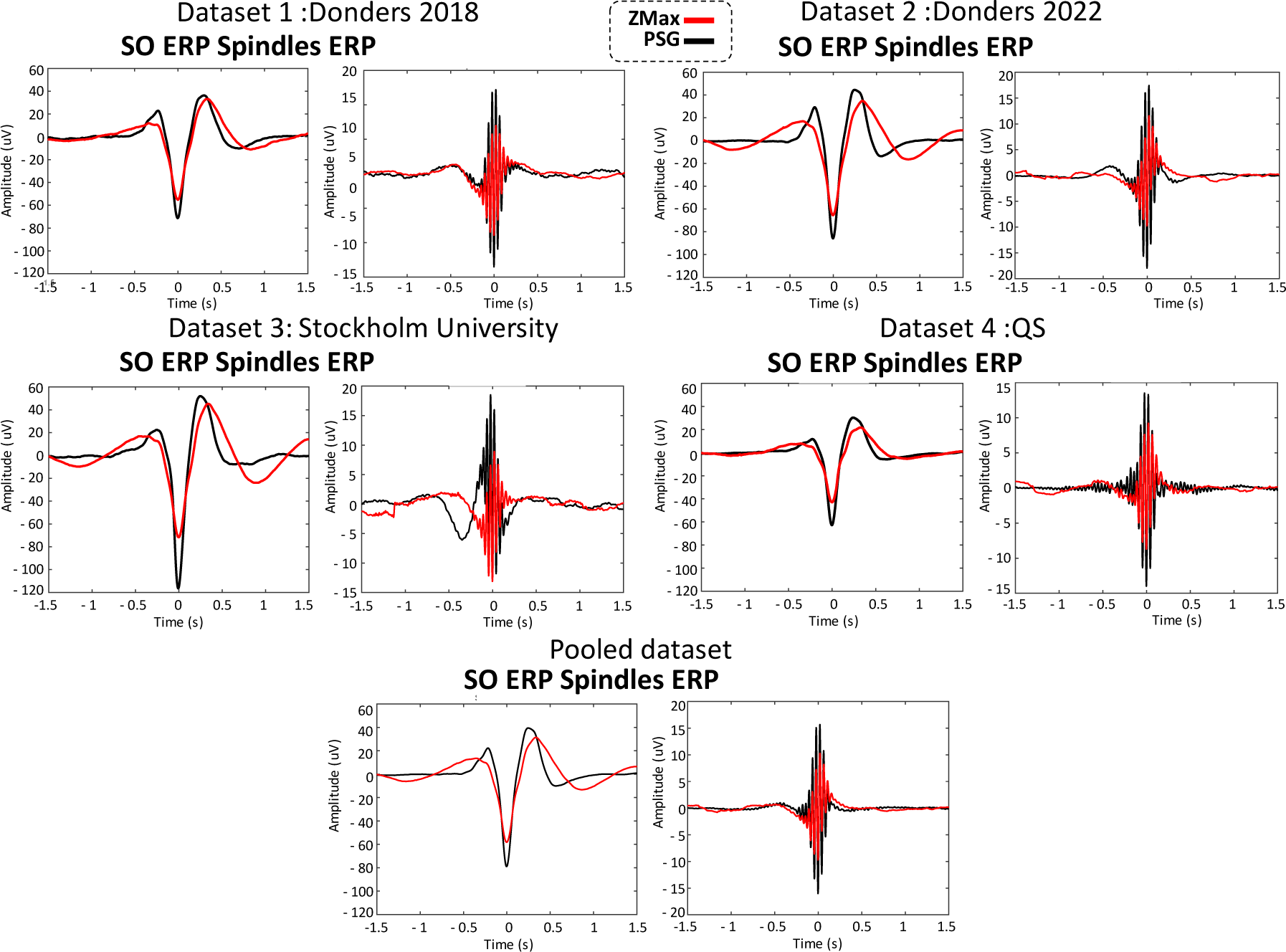
Comparison of the event-related potential (ERP) of the automatically detected slow-oscillation (SO) and spindle events in ZMax and PSG signals using SleepTrip toolbox. The results were derived by averaging the detected events over F7-Fpz and F8-Fpz for ZMax (red signal) and F3 and F4 referenced to contralateral mastoid for PSG (black). In dataset 3, due to the availability of only the F4 channel, the resulting ERP from PSG F4 were compared with the averaged ZMax ERP across F7 and F8 channels, which created a slightly different signal when compared with the other datasets.

Comparing the characteristics of non-REM and REM microstructural features (see Figure 9 and Supplemental Table 6) we observed a strong correlation between the outputs of ZMax and PSG across different datasets as well as the pooled dataset: SO counts (r=0.74, p<0.001), SO density per epoch (r=0.72, p<0.001), SO mean duration (0.41, p<0.001), SO mean frequency (r=0.44, p<0.001), SO trough-to-peak amplitude (r=0.55, p<0.001), SO zero-crossing slope (r=0.52, p<0.001), SO trough-to-peak lope (r=0.56, p<0.001), spindle counts (r=0.67, p<0.001), spindle density (r = 0.46, p<0.001), spindle duration (r=0.58, p<0.001), spindle trough-to-peak amplitude (r=0.44, p<0.001), spindle linear regression frequency slope (r=0.70, p<0.001), REM counts (r=0.66, p<0.001), and REM density (r=0.60, p<0.001). However, a few measures relating to REM events did not reach significant correlation level between the PSG and ZMax outcome, namely REM mean duration (r = 0.12, p = 0.22) and REM mean speed (r=0.15, p = 0.10).

**Figure 9.**
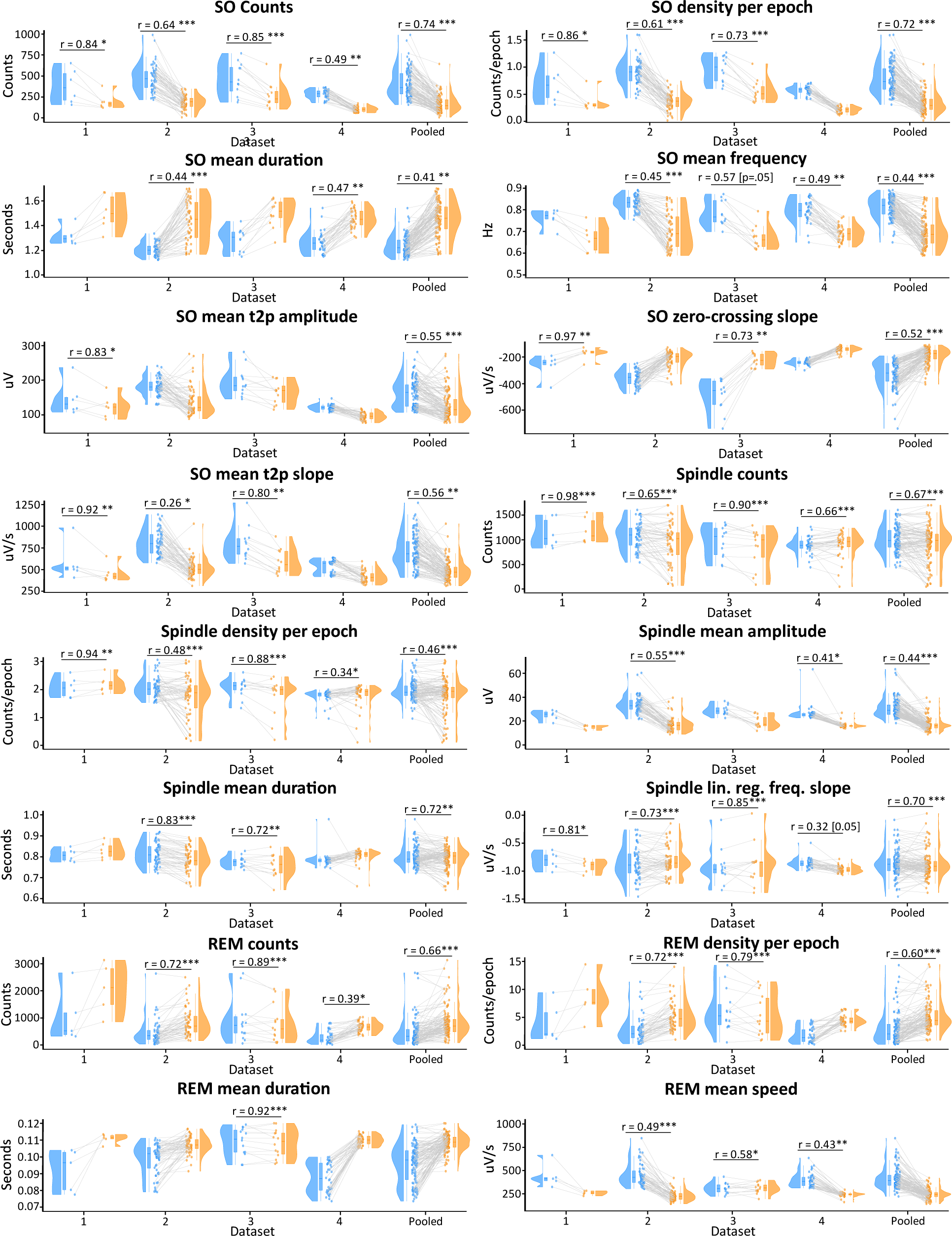
Rain-cloud plots representing the correlation between the characteristics of the microstructural features of non-REM (SO and spindles) and REM sleep (rapid eye movement events) derived from the PSG (blue) and ZMax (orange). The significant correlation values are indicated in the figure. *: p-value < 0.05, ** p-value < 0.01, ***: p-value < 0.001. T2p: trough-to-peak.

The mean difference (PSG measure vs ZMax) of sleep microstructural features was assessed using Bland-Altman plots. Our results indicate the presence of the following biases: 238.81 ± 113.91 for SO counts, 0.46 ± 0.18 counts per epoch for SO density, -0.22 ± 0.13 seconds for SO duration, 0.12 ± 0.06 Hz for SO frequency, 35.43 ± 37.26 uV for SO amplitude, -138.77 ± 82.56 uV/second for SO zero-crossing slope, 218.60 ± 153.39 uV/s for SO trough-to-peak slope, 88.72 ± 268.58 for spindle counts, 0.19 ± 0.55 counts per epoch for spindle density, 13.75 ± 6.87 uV for spindle amplitude, 0.01 ± 0.04 seconds for spindle duration, 0.02 ± 0.17 uV/second for spindle linear regression frequency slope, -364.57 ± 442.80 for REM counts, -2.16 ± 2.33 counts per epoch for REM density, -0.01 ± 0.01 seconds for REM duration, and 172.62 ± 113.44 uV/seconds for REM speed (see Figure 10).

**Figure 10.**
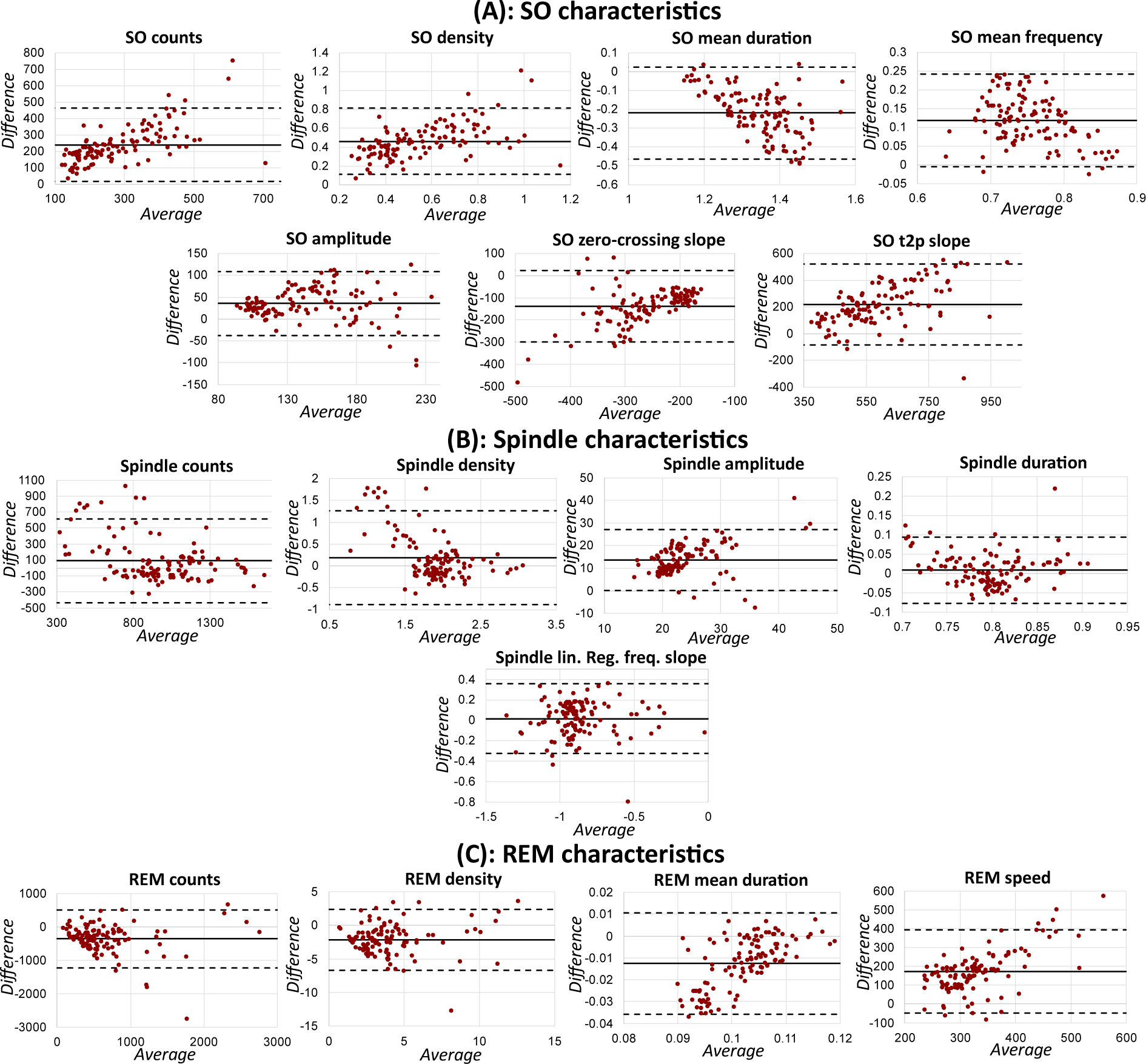
Bland-Altman plots representing the agreement of measurement of sleep microstructural features between ZMax and the PSG. **(A)** SO characteristics, **(B)** Spindle characteristics, and **(C)** REM events characteristics. The horizontal solid line in each figure represents the mean bias. The two dashed horizontal lines show the upper and lower limits of agreement. The results are demonstrated based on the pool dataset. The difference between the measures was based on PSG outcome minus the ZMax-based measure. In each plot, the x-axis represents the average of the two measures, whereas the difference between measures is shown on the y-axis.

## 3. Discussion

Due to advancements in the miniature electronics industry and the widespread implementation of AI in addition to the increased demand for health tracking, the consumer market offers an increasing number of wearable sleep sensing systems. Despite the accessibility, the extent to which wearable systems can accurately measure the intended variables is somewhat uncertain. To utilize wearables such as the ZMax EEG headband in scientific and clinical settings, it is necessary to validate their performance compared with a standard PSG system. We did this by analyzing four datasets collected by the varying research institutes in both laboratory and naturalistic home environments, resulting in a large and diverse sample. We also included a fifth dataset as a proof of concept for the applicability of ZMax headband in measuring longitudinal sleep data in clinical settings.

When considering wearable systems for data collection, the initial inquiry pertains to their feasibility of data collection. This can be evaluated across different domains such as the technology acceptance rate by the users and the rate of resulting useful data. Our behavioral analysis indicated that the ZMax sleep disturbance rate and comfort level, as well as its influence on sleep quality, morning mood, and number of awakenings, all lay within an acceptable range (see Figure 4). While we observed different sleep quality scores in a post-experimental period compared with both pre-experimental and experimental phases, it is essential to note that the extent of differences were overall small. The same applies to the reported number of awakenings. Furthermore, no compelling evidence was found to establish an association (either positive or negative) between sleep quality, morning mood, number of awakenings, ZMax disturbance, and comfort as time progressed during the experimental period. These findings suggest that the ZMax headband does not have a considerable adverse effect on sleep or mood, and that longitudinal recordings with the headband do not lead to significant deterioration of these aspects over time.

In the largest dataset of the present study (dataset 2, see Table 1), among the 37 recruited participants, we experienced only four dropouts (11% of the recruited participants) due to discomfort of the headband. Considering the extensive duration of the study, spanning 6 weeks with 15 nights of nocturnal sleep using the ZMax (including 3 nights of PSG) we deem this dropout rate to be rather moderate.

Regarding the usefulness rate of data, we observed both acceptable (e.g., 68% and 63% for datasets 2 and 4, respectively) and lower-than-expected (26% and 30% for datasets 1 and 3, respectively) rates for simultaneous ZMax-PSG recordings among healthy populations. In dataset 2, which involved our largest sample size as well as the highest rate of useful data, we provided the participants with an information brochure, as well as an instruction and troubleshooting video. Our participants were informed at their first lab visit that electrophysiological studies with EEG might result in varying degrees of skin sensitivities due to alcohol usage and chemical reactions at the wet electrode sites. Thus, we instructed the participants to clean their foreheads not too intensely, and further to adjust the headband to a tight, but not too tight fight. Furthermore, during each lab visit, the participants received feedback from the researcher regarding the quality of the collected data in addition to some strategies by which the data quality could be improved. These Goldilocks-principled instructions may have contributed to the overall acceptable tolerability of the headband even under repeated use. While we did not receive any reports on significant cases of skin irritation from participants in our studies, we also need to mention that we did not assess this aspect systematically. In dataset 4, encompassing several recordings conducted by a citizen neuroscientist, out of the 25 excluded recordings due to poor quality, 17 occurred during the last 18 recordings. This suggests the possibility of a technical issue with the wearable at the end of the study; whereas a much higher success rate was observed in the earlier phase of the study. Accordingly, with proper instruction, training, and occasional follow-ups, longitudinal home-recordings appear to be possible with considerable reliability. Of note, in datasets 1 and such instructions had not been implemented yet, which may partly explain the lower rate of successful recordings. We thus urge other researchers to adopt those instructions or similar practices when utilizing the ZMax wearable.

The analysis of sleep data is fundamentally dependent on the process of sleep scoring. Thus, when evaluating the performance of an investigational system for sleep measurement, the primary consideration should be the system’s ability to produce data that can be accurately scored. We showed the feasibility of human scoring of ZMax data, even in real-time, elsewhere (Esfahani et al., 2022b). When it comes to autoscoring, the majority of the state-of-art algorithms have been trained on standard PSG data using virtual EEG montages, e.g., mainly a frontal channel referenced to a central (e.g., Hsu et al., 2013; Supratak et al., 2020; Tsinalis et al., 2016). This is because such a montage provides information regarding rapid-/slow- eye movements through frontal channels which are useful for REM and N1 detection, and facilitates spindle detection from central scalp regions which is useful for N2 vs N3 distinction. Additionally the dominant alpha activity in the occipital regions during wake with the eyes closed may be more evident in central regions, when compared to frontal solely. Wearables such as ZMax, however, typically grant access to the frontal electrodes only, employ different electronics (e.g., amplifier) which influence the output signal, and record data with relatively higher impedance (when compared to cup electrodes from PSG) due to the use of hydrogel disposable electrodes, making these data more challenging for the conventional PSG-evaluated autoscoring algorithms. Therefore, these algorithms may not perform ideally on wearable data, if utilized ‘out of the box’ and without further fine-tuning. Alternatively, annotated data from wearables could be included into the training sets of large cross-cohort and cross-PSG system sleep staging projects like U-Sleep (Perslev et al., 2021) or YASA (Vallat et al., 2021) to make these algorithms robust to score EEG data from wearables reliably.

While our results showed that DreamentoScorer outperformed ZLab by achieving a higher recall (true positive rate among all the actual positives), precision (true positive rate among all the positive detections), and F1-score (the harmonic mean of precision and recall) across all the sleep stages as well as higher overall accuracy, Cohen’s kappa score, and macro-F1 (the average of F1-scores) across the pooled dataset, it is essential to acknowledge that due to the utilization of a cross-validation in testing DreamentoScorer performance, a direct comparison of the results derived from DreamentoScorer and ZLab may not be applicable. This is because ZLab has been trained on a completely different dataset, where the number of trained data and any further details remain undisclosed due to its proprietary nature. Nevertheless, we have indicated the possibility of developing an accurate autoscoring system for ZMax EEG data that results in relatively acceptable outcomes as detailed in Tables 2 and 3. Further research could focus on development of an artifact rejection algorithm for DreamentoScorer (in progress). This algorithm would enable DreamentoScorer to assess the data quality, rejecting artifactual epochs, and then proceed to apply autoscoring exclusively on the data segments with acceptable quality.

Considering the sleep micro- and macro-structural alterations with aging (Crowley et al., 2022; Nicolas et al., 2022; Van Cauter et al., 2000), it was found that our autoscoring algorithm, DreamentoScorer, which was trained on a relatively young population, would not work ideally for older age groups. We recommend future studies on older populations and patient groups to employ DreamentoScorer cautiously and apply a post-autoscoring step according to our data quality assessment procedure to ensure the alignment between the sleep stages and the TFR alterations (see Figure 6 and Supplemental Figure 1). Collecting simultaneous PSG-ZMax data from elderly is crucial to developing more personalized models tailored to their altered sleep stage characteristics.

An important use case of wearables for longitudinal studies is their ability to derive sleep statistics. Based on our findings, both ZLab and DreamentoScorer have relatively acceptable results while determining SPT, TST, SE, SME, SOL, WASO, etcetcra. Of note, similar to other autoscoring approaches, we have observed the issue of early REM detections in DreamentoScorer which also influenced the lower estimations of REM latency that should be considered by the users. To overcome this issue, we proposed a method to replace the REM episodes detected before the first N2 detection with N1. While this method was deactivated in evaluating datasets 1-4, we activated it to assess dataset 5. Future work may integrate innovative features as a potential solution to the problem of misidentifying N1 as early REM sleep.

Wearable systems typically utilize electronics which are relatively different from the ones used in a PSG system (e.g., amplifiers and hardware filters configuration). Despite not anticipating a strong agreement between the measured relative bandpower in using ZMax and PSG, our findings revealed that the relative bandpower calculated using ZMax generally exhibited only a minor bias. These findings imply that ZMax can effectively capture a dependable EEG signal, allowing for the accurate detection of crucial frequencies required for sleep recording within the 0.3–30 Hz range.

Beyond passive sleep assessment, another intriguing application of wearables is their capability to ‘modulate’ sleep. Sleep modulation is typically subdivided into (1) non-REM sleep modulation, e.g., to enhance deep sleep and memory consolidation using CLAS and TMR techniques, or (2) REM sleep modulation for studying (lucid) dreams. In our previous work, we have introduced Dreamento as an open-source tool that allows conducting such studies with wearables (Esfahani et al., 2022a). In this work, we have deemed to address the similarities and differences between the characteristics of the oscillations such as SO, spindles, and REM events in ZMax with respect to PSG signal which have to be considered either for sleep modulation studies or to conduct post-processing of the microstructural features of sleep. While the overall characteristics of the ZMax ERPs resemble those from PSG, there were notable discrepancies that should be acknowledged (see Figure 8 and Supplemental Table 6). The ZMax signals yielded SO and spindle ERPs with consistently lower amplitude when compared to the PSG outcome (see Table 3 for detailed results). Nevertheless, this might have been expected due to the different montage configuration of the analysis. More specifically, while the channels used for PSG event detection were more spatially separated (F3 and F4 referenced to the mastoids), ZMax F7 and F8 channels were referenced to a relatively close channel (Fpz). This difference could contribute to the fact that the detected fluctuations in ZMax signal are typically attenuated when compared to the PSG. Moreover, the ZMax SO ERPs consistently exhibited wider trough-to-peak and peak-to-trough intervals when compared to the PSG signals. Nevertheless, our analysis based on different datasets indicated a clear association between the SO and spindle events in both ZMax and PSG across all datasets (see Figures 8 & 9). Notably, in the current study, we employed identical parameters for detecting SO, spindle, and REM events in both ZMax and PSG (Supplemental Table 2). The intention behind this approach was to standardize all the factors, allowing for a direct comparison of the actual signals recorded in ZMax vs PSG. We encourage future studies to consider the addressed discrepancies that are expected to be observed in ZMax signal to either aim for a more precise real-time detection of the desired oscillation or to conduct a more accurate post-processing of the desired microstructural events. This would then subsequently enhance the validity and reliability of the findings.

## 4. Conclusions

In this study, we evaluated the performance of the sleep EEG wearable ZMax, compared to PSG as ground truth. ehaviorally, ZMax does not appear to influence its wearer’s sleep and mood negatively, and we did not observe an accumulative deteriorative effect of wearing ZMax over time. Our analysis using four different datasets with simultaneous ZMax-PSG recordings showed that ZMax provides acceptable signal quality. ZMax signals can be fed to machine-learning algorithms such as ZLab and DreamentoScorer which both resulted in reliable sleep assessment metrics. Nevertheless, DreamentoScorer was more accurate in identifying sleep stages and its corresponding metrics except for REM latency. We have evaluated the agreement between the output measures of ZMax vs PSG for different purposes, containing bandpower, non-REM microstructural features such as SO and spindles, and REM features including rapid eye movement instances. Despite the discrepancies observed in the ZMax outcomes, if future studies take careful consideration of these findings, ZMax has the potential to serve as a dependable tool for both sleep monitoring and modulation.

## Supporting information

ZmaxValidation_bioRxiv_Supplementary_Material

## Acknowledgments

This work was supported by the PPP Allowance made available by Health∼Holland, Top Sector Life Sciences & Health. The authors would like to thank Eva Spetter, and Leonie Kooloos for their contribution in data collection.

## Author Contributions

Conceptualization: MJE, FDW, MD; Data curation: MJE, FDW, MB, YK, RtH, CH, MtA; Funding acquisition: MD; Methodology/Software: MJE, FDW; Formal analysis: MJE; Investigation: MJE, FDW, MB, YK, ES, RtH, CH, MtA, MP, TS; Supervision: FDW, MD; Visualization: MJE; Writing – original draft: MJE; Writing – review & editing: MJE, FDW, MB, TA, SA, MvH, YK, ES, LB, NA, MtA, MP, RtH, CH, TS, JA, MD.

## Notes

https://github.com/dreamento/dreamento

### Competing Interest Statement

The authors have declared no competing interest.

