## Supplementary material for "Validation of the sleep EEG headband ZMax": ZmaxValidation_bioRxiv_Supplementary_Material

### Supplemental Information

#### Supplemental Figures

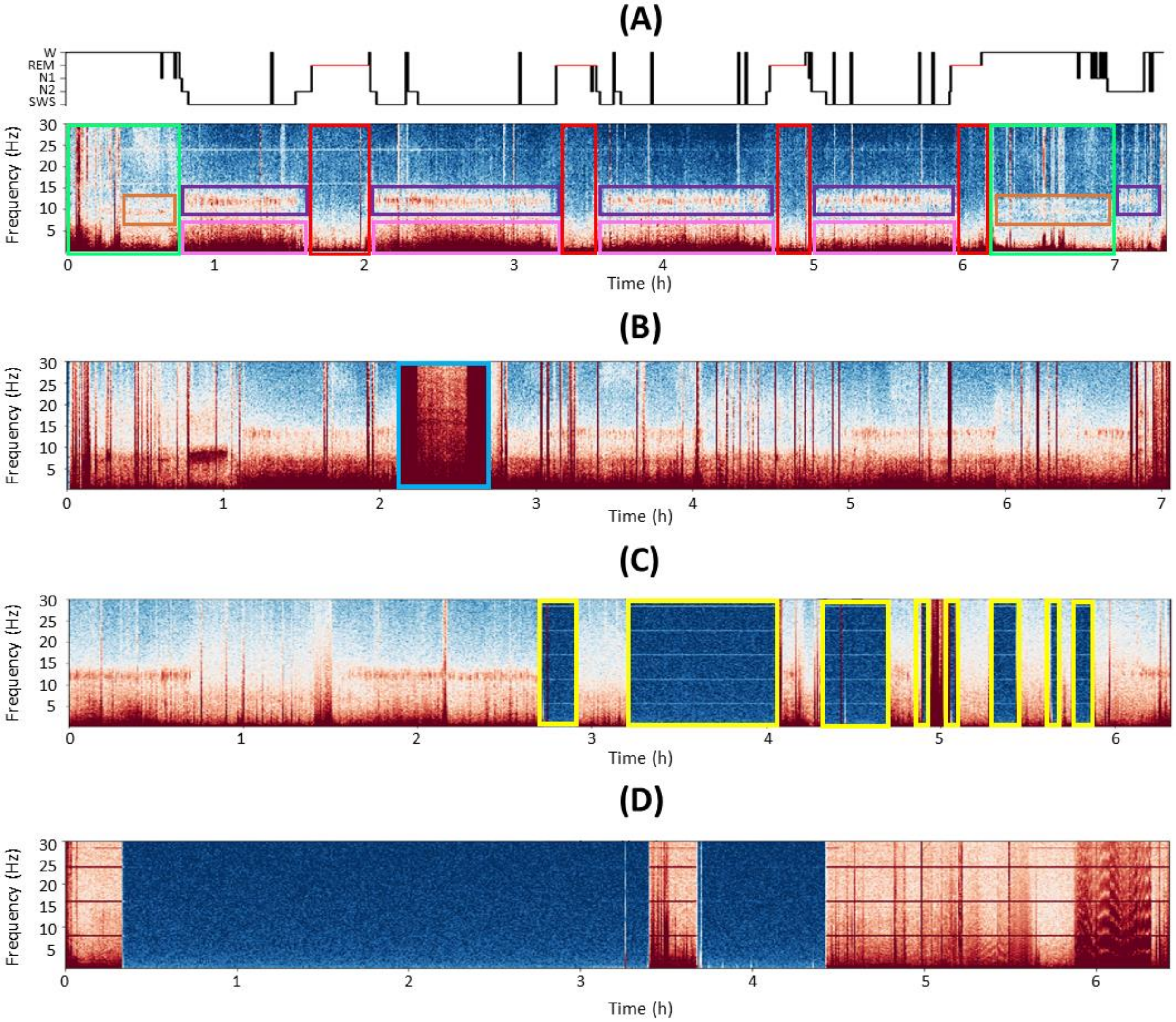

***Supplemental Figure 1.*** **EEG data quality assessment protocol using time-frequency representations (TFR).** Initially, a preliminary assessment of EEG data quality is conducted by examining the TFR of the complete data in the primary sleep frequency range of 0.1–30 Hz. This approach evaluates the clarity of sleep stage transitions through the visibility of relevant events and microstructural features upon cursory examination. **(A)** Sample TFR of a clean EEG data from a single channel and the corresponding hypnogram. High-quality EEG data is characterized by the following distinct events: (1) high power of 0.5–4 Hz SO activity (pink boxes) and 12–15 Hz spindle activity (purple boxes) as an indication of N2 and SWS sleep, (2) mixed-frequency band power as an indication of N1 and REM sleep (red boxes), and (3) the wakefulness episodes (green boxes) comprising bursts of broad-frequency activity (in case of movement) and 8–12 Hz alpha activity while being awake with the closed eyes. **(B)** Noisy epochs (blue box) characterized by high power across a broadband frequency range obscuring the identification of sleep microstructures, e.g., spindles and SOs. **(C)** Data loss or disconnection epochs (yellow box) indicated by the absence of power throughout the frequency range. This sample also represents a fragmented data after ~2.5 hours of recording. **(D)** completely useless data including long periods of fragmented data, noise, and data loss. While completely useless data should be thoroughly excluded from subsequent processing, for the remaining (depending on the goal of research), one needs to explore the time-domain data using conventional scoring software, e.g., SpiSOP, Domino to accurately mark and exclude epochs contaminated with poor data quality.

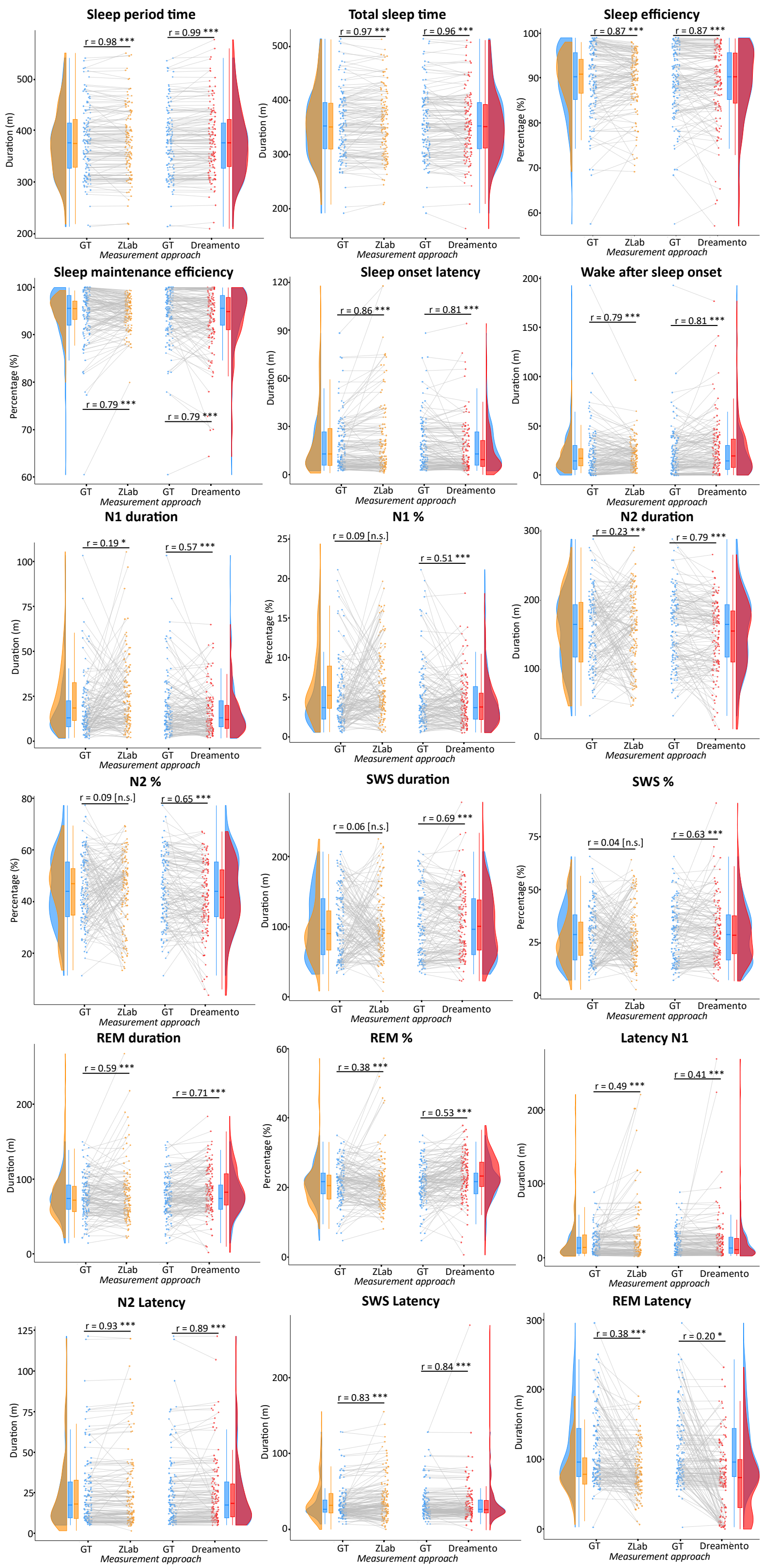

***Supplemental Figure 2. Raincloud plots representing the agreement between the ground truth sleep statistics in comparison with the results derived from ZLab and DreamentoScorer.*** *The correlation values with the corresponding p-values are represented. GT: ground truth. *: p <.05, **: p < .01, ***: p < .001. n.s.: not significant correlation.*

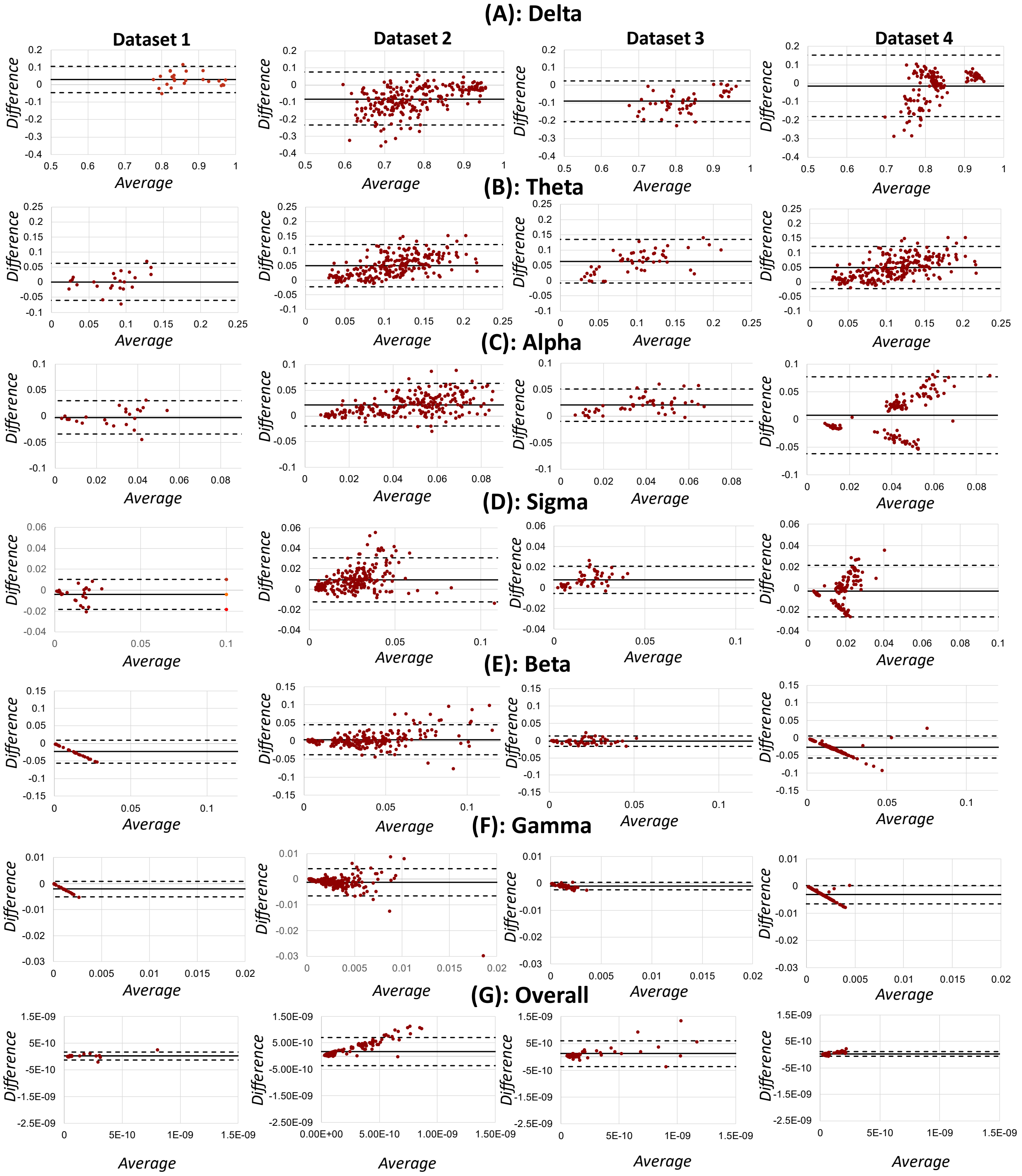

***Supplemental Figure 3. Bland-Altman plots representing the agreement between the relative bandpower calculate based on ZMax and PSG recordings for each dataset, separately.*** *(****A****) delta-band (0.5–4 Hz) power, (****B****) theta-band (4–8 Hz) power, (****C****) Alpha-band (8–12 Hz) power, (****D****) Sigma-band (12–16 Hz) power, (****E****) Beta-band (16–30 Hz) power, (****F****) Gamma-band (30–40 Hz) power, and (****G****) absolute overall power. First column: dataset 1, second column: dataset 2, third column: dataset 3, fourth column: dataset 4. The x-axis represents the average of the ZMax and PSG outcome and the y-axis demonstrated the difference between PSG and ZMax outcome. The same scales as in Figure 7 of the manuscript has been used in this figure for better comparison purposes.*

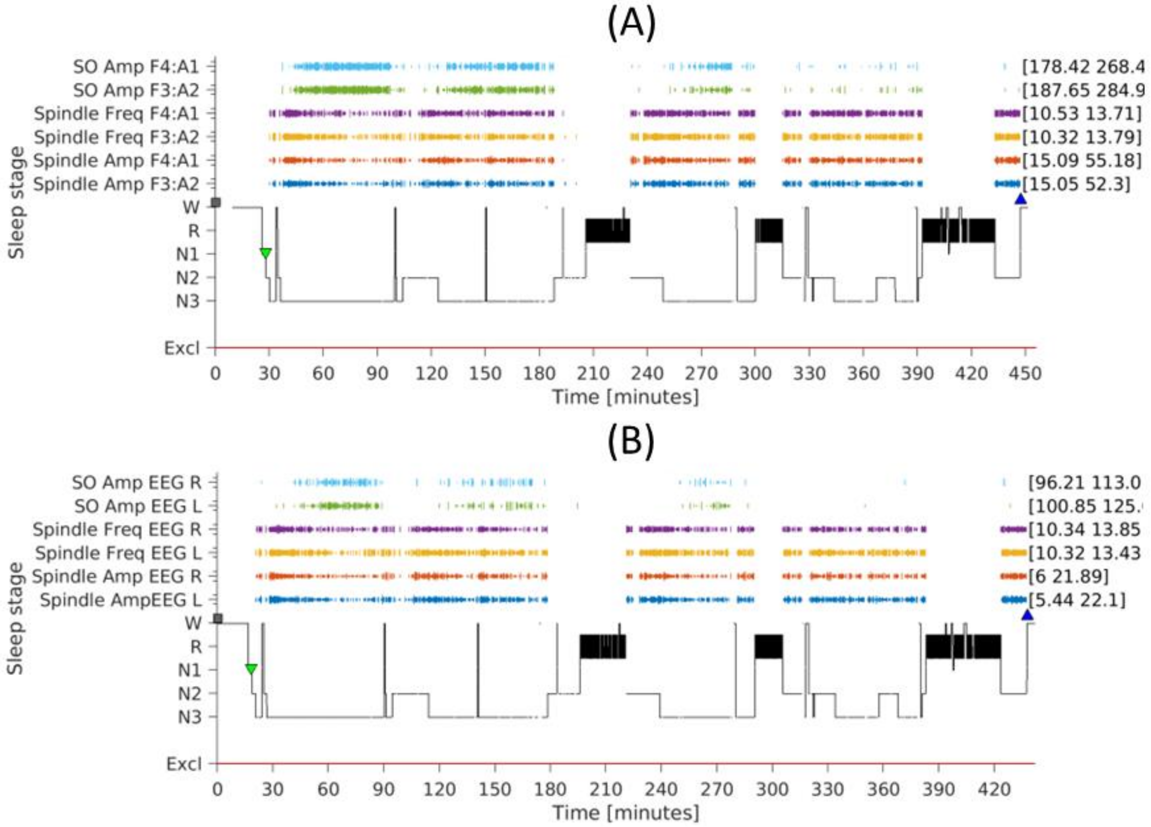

***Supplemental Figure 4. Sample hypnogram integrated with the detected SO and spindle events.*** *The detection's associated features, such as amplitude or frequency range, are indicated within brackets on the right side of the panel. Any artifact-containing epochs were excluded, and the corresponding stage is omitted from the hypnogram (indicated by no trace of hypnogram at some points).*

#### Supplemental Tables

| Parameter | Value range |
|---|---|
| boosting_type | ['gbdt'] |
| n_estimators | sp_randint(100, 5000) *(best = 2890)* |
| max_depth | sp_randint(1, 10) *(best = 6)* |
| num_leaves | sp_randint(10, 100) *(best = 64)* |
| colsample_bytree | sp_uniform(loc=0.1, scale=0.9) *(best = 0.8)* |
| importance_type | ['gain'] |
| learning_rate | sp_uniform(loc=0.01, scale=0.1) *(best = 0.03)* |
| objective | ['multiclass'] |
| class_weight: | [{0: 1.5, 1: 1.5, 2: 1, 3: 1, 4: 1}] |
| min_split_gain | sp_uniform(loc=0, scale=0.1) *(best = 0.003)* |
| min_child_weight: | sp_randint(1, 10) *(best = 3)* |
| min_child_samples | sp_randint(10, 30) *(best = 18)* |
| subsample | sp_uniform(loc=0.1, scale=0.9) *(best = 0.225)* |
| reg_alpha | sp_uniform(loc=0, scale=0.1) *(best = 0.056)* |
| reg_lambda | sp_uniform(loc=0, scale=0.1)} *(best = 0.07)* |

***Supplemental Table 1. The randomized grid search hyperparameters configuration to determine the best set of parameters for DremaentoScorer.*** *The best values were identified while running the search on the first iteration of cross-validation on dataset 2. The number of iterations for the randomized grid search was set to 10, with a 5-fold cross-validation, and a random state equal to 42.*

| *Aim* | *SleepTrip function* | *Configurations* |
|---|---|---|
| ***SO detection*** | *st_slowwaves(cfg);* | *cfg.stages = {'N2','N3'};<br>cfg.thresholdstages = cfg.stages;<br>cfg.minfreq = 0.50;<br>cfg.maxfreq = 1.11;<br>cfg.minfreqdetectfilt = .3;<br>cfg.maxfreqdetectfilt = 3.5;<br>cfg.meanfactoramp = 1.5;<br>cfg.meanfactordwnpk = 1.5;<br>cfg.minamplitude = 0;<br>cfg.maxamplitude = 600;<br>cfg.minuppeak = 10;<br>cfg.maxdownpeak = -15;<br>cfg.filterSDamp = 3;<br>cfg.filterSDdur = 3;<br>cfg.filterSDslopes = 3;* |
| ***Spindle detection*** | *st_freqpeak(cfg): detecting individuals' spindle peak frequency* | *cfg.peaknum = 1 ;<br>cfg.foilim = [6, 18];<br>cfg.smooth = .3;<br>cfg.minpeakdist = 2 ;<br>cfg.powlawnorm = 'yes';<br>cfg.pownorm = 'no';<br>cfg.prepeak = 1;<br>cfg.postpeak = 1;* |
| | *st_spindles(cfg): detecting spindle events* | *cfg.stages = {'N2','N3'};<br>cfg.thresholdstages = cfg.stages;<br>cfg.centerfrequency = freqpeak;<br>cfg.leftofcenterfreq = 1.5;<br>cfg.rightofcenterfreq = 1.5;<br>cfg.thresholdaggmeth = 'respectivechan';<br>cfg.factorthreshbeginend = 1.5;<br>cfg.envelopemeth = 'smoothedRMSwd';<br>cfg.factorthresholdcriterion = 1.5;<br>cfg.minduration = .5;<br>cfg.maxduration = 2;<br>cfg.mergewithin = 0;<br>cfg.minamplitude = 0;<br>cfg.maxamplitude = 120;<br>cfg.filterSDamp = 3;<br>cfg.filterSDdur = 3;<br>cfg.filterSDfreq = 3;<br>cfg.thresholdformbase = 'std';<br>cfg.thresholdsignal = 'filtered_signal';<br>cfg.rmstimewndw = .2;<br>cfg.movavgtimewndw = .2;* |
| ***REM detection*** | *st_rems(cfg);* | *cfg.stages = {'R' };<br>cfg.channel = {'EEG L', 'EEG R'};<br>% or {'EOG1:A1', 'EOG2:A1'};* |

***Supplemental Table 2. The configuration of SleepTrip (RRID: SCR_017318, https://github.com/Frederik-D-Weber/sleeptrip) functions to detect REM and non-REM microstructural features.*** *For SO and spindle features analysis, F3:A2 and F4:A1 channels from the PSG and F7-Fpz and F8-Fpz from ZMax were used. For REM features identification, the dedicated EOG channels from the PSG (EOG1:A1 and EOG2:A1) and the two EEG/EOG channels from ZMax (F7-Fpz and F8-Fpz) were employed.*

| | | Mean Difference | SE | t | Cohen's d | $p_{bonf}$ | $p_{holm}$ |
|---|---|---|---|---|---|---|---|
| **Post Hoc Comparisons - Sleep quality** | | | | | | | |
| Pre-experimental | Experimental | 0.142 | 0.079 | 1.804 | 0.217 | 0.228 | 0.076 |
| | Post-experimental | -0.191 | 0.079 | -2.428 | -0.291 | 0.054 | **0.036** |
| Experimental | Post-experimental | -0.333 | 0.079 | -4.233 | -0.508 | < .001 | **< .001** |
| **Post Hoc Comparisons - Morning mood** | | | | | | | |
| Pre-experimental | Experimental | 0.028 | 0.165 | 0.172 | 0.025 | 1.000 | 0.864 |
| | Post-experimental | -0.286 | 0.165 | -1.732 | -0.251 | 0.265 | 0.185 |
| Experimental | Post-experimental | -0.315 | 0.165 | -1.904 | -0.275 | 0.185 | 0.185 |
| **Post Hoc Comparisons - Number of awakenings** | | | | | | | |
| Pre-experimental | Experimental | -0.094 | 0.131 | -0.718 | -0.101 | 1.000 | 0.475 |
| | Post-experimental | 0.412 | 0.131 | 3.144 | 0.443 | 0.008 | **0.005** |
| Experimental | Post-experimental | 0.506 | 0.131 | 3.862 | 0.544 | < .001 | **< .001** |

***Supplemental Table 3. Results of repeated measures ANOVA over behavioral analysis.*** *The bolded values correspond to statistically significant p-values.*

| ***Frequency band*** | ***Mean bias*** | ***Std*** | ***LLOA*** | ***ULOA*** |
| --- | --- | --- | --- | --- |
| ***Delta*** | *-0.05531* | *0.08534* | *-0.22258* | *0.111959* |
| ***Theta*** | *0.044801* | *0.043765* | *-0.04098* | *0.130579* |
| ***Alpha*** | *0.01598676* | *0.027087375* | *-0.0371* | *0.069078* |
| ***Sigma*** | *0.00461* | *0.012293* | *-0.01948* | *0.028705* |
| ***Beta*** | *-0.00824* | *0.02265* | *-0.05263* | *0.036154* |
| ***Gamma*** | *-0.00185* | *0.00242* | *-0.00659* | *0.002893* |
| ***Overall absolute power*** | *1.13312E-10* | *2.21876E-10* | *-3.21564E-10* | *5.48189E-10* |

***Supplemental Table 4. Measures of bandpower agreement between ZMax and PSG over the pooled dataset.*** *Std: standard deviation, LLOA: lower limit of agreement, ULOA: upper limit of agreement.*

| ***Freq. band*** | ***Dataset*** | ***N1*** | | | ***N2*** | | | ***SWS*** | | | ***REM*** | | |
|---|---|---|---|---|---|---|---|---|---|---|---|---|---|
| | | *ZMax* | *PSG* | *Corr.* | *ZMax* | *PSG* | *Corr.* | *ZMax* | *PSG* | *Corr.* | *ZMax* | *PSG* | *Corr.* |
| ***Delta*** | *1* | *4.24e-11 ± 3.30e-11* | *5.18e-11 ± 4.06e-11* | *0.99 **** | *3.38e-11 ± 3.08e-11* | *8.34e-11 ± 6.08e-11* | *0.99 **** | *3.33e-10 ± 1.70e-10* | *3.72e-10 ± 2.73e-10* | *0.85 ** | *2.45e-11 ± 7.50e-12* | *3.64e-11 ± 1.30e-11* | *-0.03 [0.95]* |
| ***Theta*** | | *3.99e-12 ± 2.24e-12* | *5.84e-12 ± 3.09e-12* | *0.67 [0.14]* | *5.00e-12 ± 3.37e-12* | *7.59e-12 ± 2.88e-12* | *0.24 [0.65]* | *1.07e-11 ± 6.24e-12* | *1.01e-11 ± 3.58e-12* | *0.03 [0.95]* | *3.09e-12 ± 8.37e-13* | *3.89e-12 ± 2.26e-12* | *0.63 [0.18]* |
| ***Alpha*** | | *1.32e-12 ± 4.73e-13* | *2.21e-12 ± 5.83e-13* | *0.67 [0.14]* | *1.84e-12 ± 4.60e-13* | *1.45e-12 ± 3.46e-13* | *0.29 [0.58]* | *2.66e-12 ± 5.47e-13* | *1.12e-12 ± 2.52e-13* | *0.27 [0.60]* | *9.18e-13 ± 2.58e-13* | *1.51e-12 ± 4.82e-13* | *0.87 ** |
| ***Sigma*** | | *7.38e-13 ± 2.24e-13* | *9.08e-13 ± 1.63e-13* | *0.38 [0.45]* | *8.82e-13 ± 1.60e-13* | *5.47e-13 ± 6.51e-14* | *0.76 [0.08]* | *8.89e-13 ± 1.32e-13* | *4.78e-13 ± 9.38e-14* | *0.94 *** | *4.94e-13 ± 1.41e-13* | *5.69e-13 ± 9.54e-14* | *0.44 [0.38]* |
| ***Beta*** | | *1.41e-12 ± 2.20e-13* | *6.17e-14 ± 8.84e-15* | *-0.36 [0.47]* | *9.19e-13 ± 2.87e-13* | *3.99e-14 ± 4.15e-15* | *-0.09 [0.87]* | *7.95e-13 ± 3.19e-13* | *4.31e-14 ± 1.22e-14* | *-0.23 [0.67]* | *1.02e-12 ± 2.59e-13* | *3.66e-14 ± 4.90e-15* | *0.30 [0.57]* |
| ***Gamma*** | | *1.29e-13 ± 1.35e-14* | *3.68e-16 ± 1.56e-16* | *0.68 [0.13]* | *7.44e-14 ± 1.49e-14* | *5.83e-16 ± 4.29e-16* | *-0.30 [0.56]* | *6.64e-14 ± 1.17e-14* | *2.90e-15 ± 2.20e-15* | *-0.28 [0.59]* | *8.73e-14 ± 1.91e-14* | *2.42e-16 ± 5.81e-17* | *0.02 [0.98]* |
| ***Delta*** | *2* | *3.39e-11 ± 2.44e-11* | *5.99e-11 ± 2.96e-11* | *0.57 **** | *2.62e-11±1.11e-11* | *1.09e-10 ± 3.30e-11* | *0.28 ** | *2.15e-10 ± 3.29e-10* | *6.21e-10 ± 2.61e-10* | *0.05 [0.71]* | *2.12e-11 ± 1.12e-11* | *6.22e-11 ± 2.02e-12* | *0.33 *** |
| ***Theta*** | | *2.97e-12 ±1.14e-12* | *1.55e-11 ± 7.00e-12* | *0.59 **** | *3.23e-12 ± 1.05-12* | *1.98e-11 ± 7.18e-12* | *0.69 **** | *8.55e-12 ± 3.18e-12* | *4.09e-11 ± 1.64e-11* | *0.76 **** | *2.66e-12 ± 9.60e-13* | *1.65e-11 ± 7.21e-12* | *0.66 **** |
| ***Alpha*** | | *1.29e-12 ± 5.21e-13* | *7.03e-12 ± 3.10e-12* | *0.58 **** | *1.70-e12 ± 8.01e-13* | *9.13e-12 ± 3.66e-12* | *0.83 **** | *2.98e-12 ± 1.44e-12* | *1.59e-11 ± 7.90e-12* | *0.85 **** | *1.01e-12 ± 4.15e-13* | *6.22e-12 ± 2.77e-12* | *0.73 **** |
| ***Sigma*** | | *7.56e-13 ± 2.91e-13* | *3.70e-12 ± 1.36e-12* | *0.61 **** | *1.12e-12 ± 6.78-13* | *6.26e-12 ± 3.14e-12* | *0.90 **** | *1.38e-12 ± 8.68e-13* | *6.71e-12 ± 3.35e-12* | *0.87 **** | *5.34e-113 ± 2.05e-13* | *2.60e-12 ± 9.72e-13* | *0.56 **** |
| ***Beta*** | | *1.69e-12 ± 8.15e-13* | *5.71e-12 ± 2.22e-12* | *0.48 **** | *1.05-12 ± 4.51-13* | *3.83e-12 ± 1.13e-12* | *0.36 *** | *1.11e-12 ± 5.29e-13* | *3.16e-12 ± 8.75e-13* | *0.23 [0.07]* | *1.18e-12 ± 5.25e-13* | *4.31e-12 ± 1.94e-12* | *0.71 **** |
| ***Gamma*** | | *1.93e-13 ± 1.43e-13* | *3.74e-13 ± 2.37e-13* | *0.23 [0.07]* | *9.88e-14 ±5.44e-14* | *2.08e-13 ± 1.06e-13* | *0.42 *** | *1.17e-13 ± 7.22e-14* | *1.86e-13 ± 7.36e-14* | *0.29 ** | *1.09e-13 ± 5.73e-14* | *2.25e-13 ± 1.21e-13* | *0.48 **** |
| ***Delta*** | *3* | *8.15e-11 ± 6.03e-12* | *8.62e-11 ± 3.45e-11* | *0.01 [0.97]* | *8.30e-11 ± 4.91e-11* | *1.42e-10 ± 5.77e-11* | *0.24 [0.42]* | *4.85e-10 ± 3.02e-10* | *7.67e-10 ± 4.00e-11* | *0.56 ** | *5.47e-11 ± 2.82e-11* | *6.51e-11 ± 1.87e-11* | *0.23 [0.44]* |
| ***Theta*** | | *5.46e-11 ± 2.65e-12* | *2.06e-11 ± 1.59e-11* | *0.78 *** | *7.07e-12 ± 3.92e-12* | *2.42e-11 ± 1.35e-11* | *0.75 *** | *1.57e-11 ± 7.38e-12* | *4.38e-11 ± 1.94e-11* | *0.60 ** | *4.96e-12 ± 2.21e-12* | *1.72e-11 ± 9.34e-12* | *0.55 [0.05]* |

| | | | | | | | | | | | | | |
|---|---|---|---|---|---|---|---|---|---|---|---|---|---|
| ***Alpha*** | | 2.38e-12 ± 9.19e-13 | 7.35e-12 ± 3.96e-12 | 0.72 ** | 3.07e-12 ±1.60e-12 | 8.56e-12 ± 3.84e-12 | 0.70 ** | 5.00e-12 ± 7.76e-13 | 1.37e-11 ± 4.64e-12 | 0.79 ** | 1.80e-12 ± 7.29e-13 | 5.95e-12 ± 2.38e-12 | 0.59 * |
| ***Sigma*** | | 1.26e-12 ± 5.10e-13 | 3.49e-12 ± 1.74e-12 | 0.58 * | 1.61e-12 ± 6.81e-13 | 4.53e-12 ± 1.74e-12 | 0.68 ** | 1.95e-12 ± 7.76e-13 | 4.64e-12 ± 1.963-12 | 0.47 [0.10] | 8.64e-13 ± 3.47e-13 | 2.14e-12 ± 6.76e-13 | 0.40 [0.17] |
| ***Beta*** | | 2.20e-12 ± 9.07e-13 | 3.26e-12 ± 8.87e-13 | 0.63 * | 1.56e-12 ± 6.01e-13 | 1.96e-12 ± 3.69e-13 | 0.19 [0.53] | 1.86e-12 ± 6.79e-13 | 1.48e-12 ± 3.63e-13 | -0.10 [0.76] | 1.88e-12 ± 9.15e-13 | 2.77e-12 ± 8.91e-13 | 0.58 * |
| ***Gamma*** | | 2.00e-13 ± 8.35e-14 | 1.35e-13 ± 4.34e-14 | 0.90 *** | 1.30e-13 ± 5.45e-14 | 7.91e-14 ± 2.14e-14 | 0.48 [0.10] | 1.70e-13 ± 6.75e-14 | 6.86e-14 ± 1.75e-14 | 0.28 [0.36] | 1.66e-13 ± 8.83e-14 | 1.05e-13 ± 4.28e-14 | 0.90 *** |
| ***Delta*** | *4* | 3.11e-12 ± 7.91e-13 | 2.69e-11 ± 7.33e-12 | 0.25 [0.13] | 3.24e-11 ± 6.98e-12 | 6.20e-11 ± 1.01e-11 | 0.15 [0.39] | 1.136e-10 ± 2.12e-11 | 2.11e-10 ± 2.97e-11 | 0.51 ** | 3.05e-11 ± 9.51e-12 | 3.41e-11 ±6.00e-12 | 0.05 [0.78] |
| ***Theta*** | | 4.31e-11 ± 1.62e-11 | 6.84e-12 ± 2.98e-12 | 0.31 [0.06] | 4.21e-12 ± 6.63e-13 | 6.91e-13 ± 6.55e-13 | 0.19 [0.26] | 6.79e-12 ± 7.83e-13 | 9.97e-12 ± 2.55e-12 | -0.05 [0.78] | 2.95e-12 ± 6.37e-13 | 4.28e-12 + 1.26e-12 | 0.08 [0.61] |
| ***Alpha*** | | 1.44e-12 ± 3.74e-13 | 3.10e-12 ± 1.08e-12 | 0.18 [0.28] | 2.50e-12 ± 3.48e-13 | 2.06e-12 ± 1.01e-12 | 0.06 [0.74] | 2.23e-12 ± 2.34e-13 | 1.43e-12 ± 1.09e-12 | -0.09 [0.59] | 1.05e-12 ± 2.28e-13 | 2.31e-12 ± 4.66e-13 | 0.22 [0.19] |
| ***Sigma*** | | 8.32e-13 ± 1.75e-13 | 1.12e-12 ± 4.14e-13 | 0.00 [0.99] | 1.07e-12 ± 1.56e-13 | 6.91e-13 ± 6.55e-13 | 0.03 [0.87] | 8.31e-13 ± 5.04e-13 | 5.04e-13 ± 4.62 e-13 | -0.01 [0.94] | 6.31e-13 ± 1.37e-13 | 7.60e-13 ± 2.48e-13 | 0.15 [0.37] |
| ***Beta*** | | 1.69e-12 ± 5.94e-13 | 1.77e-13 ± 6.50e-13 | 0.44 ** | 1.13e-12 ± 5.27e-13 | 1.13e-13 ± 4.42e-13 | 0.15 [0.38] | 1.13e-12 ± 9.5e-13 | 8.03e-14 ± 2.73e-13 | 0.07 [0.67] | 1.42e-12 ± 5.09e-13 | 1.35e-13 ± 5.64e-13 | 0.09 [0.58] |
| ***Gamma*** | | 1.99e-13 ± 7.23e-14 | 5.74e-15 ± 33.34e-14 | –0.15 [0.36] | 1.31e-13 ± 5.50e-14 | 4.13e-15 ± 2.17e-14 | -0.06 [0.71] | 1.49e-13 ± 6.94e-14 | 4.06e-15 ± 1.53e-14 | 0.06 [0.72] | 1.48e-13 ± 4.30e-15 | 5.45e-14 ± 2.49e-14 | -0.12 [0.48] |

***Supplemental Table 5. relative (Welch) band power averaged over frontal channels.*** *The power of each ZMax and PSG signals were computed using YASA toolbox (Vallat et al., 2021). The absolute power in addition to relative power (in parentheses) of each band within different sleep stages are given in each cell. The correlation values were computed based on absolute values.*

| Event | Measure | System | Dataset 1 | Dataset 2 | Dataset 3 | Dataset 4 | Pooled dataset |
|---|---|---|---|---|---|---|---|
| SO | N recordings | | 6 | 61 | **12** | 38 | 117 |
| | Counts | PSG | 378.83 ± 186.73 | 471.97 ± 155.39 | **460.17 ± 186.47** | 282.46 ± 51.75 | 404.43 ±162.40 |
| | | ZMax | 192.92 ± 88.62 | 183.75 ± 77.99 | 263.04± 142.67 | 101.46 ± 29.48 | 165.62 ± 92.00 |
| | | Corr [p-val] | 0.84 * | 0.64 *** | 0.85 *** | 0.49 ** | 0.74 *** |
| | Density per epoch | PSG | 0.69 ± 0.32 | 0.90± 0.23 | 0.96 ± 0.23 | 0.56 ± 0.07 | 0.79 ± 0.26 |
| | | ZMax | 0.36 ± 0.17 | 0.35 ± 0.15 | 0.54 ± 0.20 | 0.20 ± 0.05 | 0.32 ± 0.17 |
| | | Corr [p-val] | 0.86 * | 0.61 *** | 0.73 ** | 0.20 [0.22] | 0.72 *** |
| | Mean duration (s) | PSG | 1.31 ± 0.07 | 1.21 ± 0.05 | 1.28 ± 0.09 | 1.26 ± 0.08 | 1.24 ± 0.08 |
| | | ZMax | 1.51 ± 0.13 | 1.44 ± 0.16 | 1.50 ± 0.10 | 1.46 ± 0.07 | 1.45 ± 0.13 |
| | | Corr [p-val] | 0.54 [0.27] | 0.44 *** | 0.53 [0.08] | 0.47** | 0.41 *** |
| | Mean frequency by duration (Hz) | PSG | 0.76 ± 0.04 | 0.83 ± 0.03 | 0.78 ± 0.05 | 0.80 ± 0.05 | 0.81 ± 0.05 |
| | | ZMax | 0.67 ± 0.06 | 0.70 ± 0.08 | 0.67 ± 0.05 | 0.69 ± 0.03 | 0.69 ± 0.07 |
| | | Corr [p-val] | 0.51 [0.30] | 0.45 *** | 0.57 [0.05] | 0.49 ** | 0.44 *** |
| | Mean t2p amplitude | PSG | 146.06 ± 43.54 | 182.13± 25.13 | 195.69 ± 39.26 | 121.46 ± 8.03 | 161.96 ± 38.47 |
| | | ZMax | 123.75 ± 29.70 | 138.28 ± 40.40 | 164.34 ± 31.35 | 96.18 ± 10.88 | 126.53 ± 39.41 |
| | | Corr [p-val] | 0.83 * | 0.10 [0.46] | 0.52 [0.08] | 0.01 [0.95] | 0.55 *** |
| | Mean slope zero crossing | PSG | -265.36 ± 77.36 | -359.77 ± 56.49 | -485.46 ± 116.61 | -242.16 ± 19.04 | -329.62 ± 95.62 |
| | | ZMax | -174.50 ± 40.08 | -214.33 ± 57.57 | -224.52 ± 46.95 | -145.11 ± 20.97 | -190.86 ± 57.24 |
| | | Corr [p-val] | 0.97 ** | -0.01 [0.93] | 0.73 ** | 0.22 [0.19] | 0.52 *** |
| | Mean slope t2p | PSG | 574.08 ± 186.89 | 807.83 ± 140.63 | 799.71 ± 184.06 | 538.36 ± 76.95 | 707.49 ± 183.92 |
| | | ZMax | 454.85 ± 94.12 | 515.38 ± 106.90 | 593.50 ± 134.50 | 418.70 ± 65.67 | 488.89 ± 113.21 |
| | | Corr [p-val] | 0.92 ** | 0.26 * | 0.80 ** | 0.15 [0.36] | 0.56 *** |
| Spindle | N recordings | | 6 | 61 | 13 | 38 | 118 |
| | Counts | PSG | 1127.75 ± 276.65 | 1079.23 ± 260.52 | 968.31 ± 301.31 | 890.5 ± 158.91 | 1008.7 ± 255.11 |
| | | ZMax | 1184.42 ± 243.25 | 918.26 ± 406.84 | 835.04 ± 381.74 | 910.03 ± 241.71 | 919.97 ± 357.53 |
| | | Corr [p-val] | 0.98 *** | 0.65 *** | 0.90 *** | 0.66*** | 0.67 *** |
| | Density per epoch | PSG | 2.08 ± 0.33 | 2.07 ± 0.36 | 2.07 ± 0.33 | 1.78 ± 0.20 | 1.97 ± 0.34 |
| | | ZMax | 2.19 ± 0.28 | 1.74 ± 0.71 | 1.76 ± 0.66 | 1.81 ± 0.41 | 1.79 ± 0.61 |
| | | Corr [p-val] | 0.94 ** | 0.48 *** | 0.88 *** | 0.34* | 0.46 *** |
| | Mean duration | PSG | 0.81 ± 0.03 | 0.81 ± 0.05 | 0.78 ± 0.02 | 0.79 ± 0.03 | 0.80 ± 0.05 |
| | | ZMax | 0.83 ± 0.04 | 0.79 ± 0.05 | 0.76 ± 0.05 | 0.81 ± 0.03 | 0.79 ± 0.05 |
| | | Corr [p-val] | 0.73 [0.10] | 0.83 *** | 0.72 ** | -0.09 [0.59] | 0.58 *** |

| | | | | | | | |
|---|---|---|---|---|---|---|---|
| | Mean t2p amplitude | PSG | 24.97 ± 3.64 | 34.13 ± 7.11 | 29.24 ± 3.96 | 26.58 ± 6.40 | 30.69 ± 7.43 |
| | | ZMax | 14.99 ± 1.71 | 16.97 ± 5.97 | 19.40 ± 4.87 | 16.38 ± 2.20 | 16.95 ± 4.87 |
| | | Corr [p-val] | 0.66 [0.15] | 0.55 *** | 0.45 [0.11] | 0.41* | 0.44 *** |
| | Linear reg. freq. slope | PSG | -0.81 ± 0.14 | -0.86 ± 0.27 | -0.92 ± 0.30 | -0.86 ± 0.08 | -0.86 ± 0.23 |
| | | ZMax | -0.91 ± 0.10 | -0.82 ± 0.24 | -0.87 ± 0.35 | -0.97 ± 0.06 | -0.88 ± 0.22 |
| | | Corr [p-val] | 0.81 * | 0.73 *** | 0.85 *** | 0.32 [0.05] | 0.70 *** |
| REM | N recordings | | 5 | 56 | 14 | 40 | 115 |
| | Counts | PSG | 1017.2 ± 883.73 | 452.55 ± 437.73 | 881.07 ± 755.44 | 273.55 ± 226.54 | 467.01 ± 511.6 |
| | | ZMax | 2089.8 ± 840.22 | 851.54 ± 513.63 | 769.57 ± 654.36 | 668.05 ± 183.13 | 831.57 ± 549.31 |
| | | Corr [p-val] | 0.28 [0.65] | 0.72 *** | 0.89 *** | 0.39 * | 0.66 *** |
| | Density per epoch | PSG | 4.04 ± 3.17 | 2.78 ± 2.33 | 6.18 ± 3.66 | 1.8 ± 1.34 | 2.90 ± 2.67 |
| | | ZMax | 8.55 ± 3.67 | 5.11 ± 2.48 | 5.34 ± 3.59 | 4.48 ± 0.98 | 5.07 ± 2.48 |
| | | Corr [p-val] | 0.09 [0.88] | 0.72 *** | 0.79 *** | 0.26 [0.10] | 0.60 *** |
| | Mean duration | PSG | 0.09 ± 0.01 | 0.1 ± 0.01 | 0.11 ± 0.01 | 0.09 ± 0.01 | 0.1 ± 0.01 |
| | | ZMax | 0.11 ± 0.0 | 0.11 ± 0.0 | 0.11 ± 0.01 | 0.11 ± 0.0 | 0.11 ± 0.0 |
| | | Corr [p-val] | -0.2 [0.74] | 0.22 [0.10] | 0.92 *** | 0.20 [0.22] | 0.12 [0.22] |
| | Mean speed | PSG | 442.69 ± 116.44 | 453.73 ± 117.35 | 309.81 ± 54.2 | 397.19 ± 75.24 | 416.07 ± 108.82 |
| | | ZMax | 259.19 ± 20.83 | 226.68 ± 53.37 | 311.98 ± 54.22 | 240.97 ± 21.05 | 243.45 ± 51.37 |
| | | Corr [p-val] | 0.70 [0.19] | 0.49 *** | 0.58 * | 0.43 ** | 0.15 [0.1] |

***Supplemental Table 6 - Comparison between the characteristics of the microstructural features of non-REM sleep between ZMax and PSG.*** *The analysis was conducted using SleepTrip toolbox and the resulting values for each measure were averaged across channels of the same device, i.e, F7-Fpz and F8-Fpz for ZMax and F3 and F4 referenced to controlateral mastoid for PSG (in dataset 3, only the F4 channel was available, so the corresponding value was computed solely based on the F4 channel).*
